# The host phylogeny determines viral infectivity and replication across *Staphylococcus* host species

**DOI:** 10.1101/2022.11.30.518513

**Authors:** Sarah K Walsh, Ryan M Imrie, Marta Matuszewska, Gavin K Paterson, Lucy A Weinert, Jarrod D Hadfield, Angus Buckling, Ben Longdon

## Abstract

Genetic similarity between eukaryotic host species is an important determinant of the outcome of virus host shifts, where a pathogen infects a novel host species. However, it is less clear if this is the case for prokaryotes where anti-virus defences can be transmitted by horizontal gene transfer and evolve rapidly. Understanding the patterns and determinants of cross-species transmissions may provide insights into the processes underlying pathogen emergence. Here, we measure the susceptibility of 64 strains of *Staphylococcus* bacteria (48 strains of *S. aureus* and 16 non-*aureus* species) to the bacteriophage ISP, which is currently under investigation for use in phage therapy. Using three methods – plaque assays, optical density (OD) assays, and quantitative (q)PCR – we find that the host phylogeny explains a large proportion of the variation in susceptibility to ISP across the host panel. These patterns were consistent in models of only *S. aureus* strains and models with a single representative from each *Staphylococcus* species, suggesting that these phylogenetic effects are conserved both within and among host species. We find positive correlations between susceptibility assessed using a binary measure of plaque assay, OD, and qPCR, but not between the continuous component of plaque assay and any other method, suggesting that plaque assays alone may be inadequate to assess host range. Together, our results demonstrate the ability of bacterial host evolutionary relatedness to explain differences in susceptibility to phage infection, with implications for the development of ISP both as a phage therapy treatment and as an experimental system for the study of virus host shifts.

## Introduction

Host shifts, where a pathogen jumps into a novel host species and establishes onward transmission, are a major source of emerging infectious diseases. While host shifts can occur with many types of pathogen, viruses are the most prolific (1,2). Consequently, viruses originating in non-human animals make up the majority of recently emerged human infections (3–5), including several human pandemics: HIV-1, which jumped into humans from chimpanzees (3,6); influenza A viruses, which commonly emerge from wild aquatic birds (7–10); and most recently SARS-CoV-2, which most likely transmitted into humans from a bat reservoir (11–13). Given the scale and speed at which emerging viruses can impact host populations, understanding the underlying causes of virus host shifts has become a major goal of infectious disease research.

The evolutionary relationships between hosts are a key factor in determining the success of a pathogen following transmission to a novel host species, and several studies have investigated the ability of host evolutionary relatedness to explain variation in infection traits. These studies have shown that virulence tends to increase (14–18), and transmission rate (14,19) and pathogen load (18,20,21) decrease with greater evolutionary distance between donor and recipient host species. These ‘distance effects’ have been seen in viruses (19–21), bacterial pathogens (22–24), fungi (25,26), and nematodes (18), as well as in reconstructions of virus host shifts and cross species transmissions in nature (19,27). Additionally, closely related species may share similar levels of susceptibility independent of evolutionary distance to the natural host. These ‘clade effects’ create a patchwork of host clades across a phylogeny that vary in their susceptibility to a pathogen, and have been demonstrated in experimental infections of fruit flies (17,21) and in pathogens that repeatedly jump between distantly related hosts in nature (28–32). These effects can act concurrently to influence host shifts, with distance effects making successful pathogen shifts into closely related hosts more likely, and clade effects allowing for host shifts into more distantly related clades of susceptible hosts.

While most studies investigate the role of eukaryotic host phylogenies, few have examined the influence of the host phylogeny on susceptibility to infection in a prokaryotic system. Host phylogenetic effects follow the conventional wisdom that more closely related species share more similar phenotypes, and so present similar environments to invading pathogens. In bacteria, horizontal gene transfer has been shown to be an important mechanism for the acquisition of genes that confer resistance to bacteriophage (viruses that infect bacteria; hereafter ‘phage’) (33). It was previously demonstrated that the transfer of plasmids between bacterial hosts is more likely between recipients with near identical genomes to the plasmid donor, but the likelihood of plasmid transfer was not correlated with genetic distance between donors and recipients at larger evolutionary distances (34). As such, mobile genetic elements containing phage resistance genes may transmit between bacterial hosts in a way that does not segregate phylogenetically. Additionally, it has been demonstrated that evolution of the bacterial core genome is almost entirely driven by recombination, and thus more determined by the spatial structure of microbes rather than clonal inheritance (35). This may lead to weaker phylogenetic signal in virus susceptibility in bacterial hosts than that seen in animals where immunity is largely vertically transmitted.

Recent years have seen a resurgence in interest in the use of phage to treat bacterial infections due to the continuing emergence of antimicrobial resistance (36–39). Phage present a promising alternative to traditional antimicrobials in that they are self-amplifying, self-limiting, and have proved effective in the treatment of drug-resistant bacterial infections such as methicillin-resistant *Staphylococcus aureus* (MRSA) (40–42) and *Pseudomonas aeruginosa* (42). The host range of phages – and so their spectrum of therapeutic efficacy – can vary from single host species (43–45) to broad host range generalists (46–48), although a consensus has yet to be reached on the most effective method for quantifying phage host range (49–51). Nevertheless, the ability to explain variation in phage susceptibility across bacterial hosts would likely be beneficial in the design of future therapies.

Here, we use a broad host range bacteriophage (Intravenous Staphylococcal Phage; ISP) and a panel of 64 *Staphylococcus* isolates (encompassing 17 *Staphylococcus* species) to investigate how patterns of bacterial susceptibility are influenced by the evolutionary relationships between hosts. ISP, a double stranded DNA virus in the family *Myoviridae*, is closely related to *Staphylococcus* phage G1 (52), and has a broad experimental host range within *S. aureus*, infecting 86% of tested isolates in a previous study (53). However, in the same study, ISP was unable to experimentally infect *S. haemolyticus*, suggesting that this host range may be species-specific. Staphylococci are a well-established model for bacteria-phage interactions, with investigations into staphylococcal phages occurring since the 1910s. Lytic staphylococcal phages have demonstrated broad host ranges (54–57), antibiofilm activity (54,58), and vary in their efficacies against bacterial infection (59–61). Several mechanisms that are important for the interaction between staphylococci and their phage have been characterised, including common cell surface receptors used for attachment (62–67), and host resistance mechanisms such as the overproduction of surface proteins to block adsorption (68–70), restriction modification systems (71–73), and CRISPR targeted degradation of phage DNA (74–77). Despite this, much remains to be understood about the interactions between phages and their hosts. An improved understanding of the mechanisms that underpin bacteria-phage interactions may provide more general insights into virus-host interactions and their implications in the emergence of viral pathogens into novel populations.

In this study we demonstrate that ISP is a broad host range bacteriophage, able to infect both within and among species of *Staphylococcus*. Given that different methods of quantifying host range can give different estimations, we assessed the susceptibility of our *Staphylococcus* panel to ISP using three methods: plaque assays, optical density assays, and quantitative PCR. We observed considerable variation in the susceptibility of the *Staphylococcus* panel to ISP and showed that a high proportion of the observed variation in susceptibility to infection was attributable to the relationship between host species.

## Methods

### *Staphylococcus* isolates

This study made use of 64 strains of *Staphylococcus*, representing a broad phylogenetic and geographic range, and spanning 17 species that were estimated to have a shared a common ancestor ∼122mya (78) (Table S1). Each sample of *Staphylococcus* was streaked on an LB agar plate and left overnight at 37 °C. A single colony was isolated and used to inoculate 5mL of LB broth, which was incubated at 37°C, 180rpm overnight. All isolates were stored in 25% glycerol at -80°C. When required, isolates were grown up by inoculating 5mL of LB broth with a scraping of the frozen culture and incubating at 37°C, 180rpm overnight. To ensure that any observed differences in susceptibility to ISP were not a function of host availability (i.e., differences in the concentration of bacterial hosts in solution), calibration curves using optical density (OD) and colony forming units (CFU)/mL were generated for each strain. Overnight cultures were then normalised by dilution in LB broth to give similar host densities prior to their use in susceptibility assays.

### Phage preparation

An isolate of ISP was kindly provided by Jean-Paul Pirnay and Maya Merabishvili at the Queen Astrid Military Hospital (Brussels, Belgium). ISP was propagated on the *S. aureus* strain 13S44S9 and extracted using chloroform: 1mL aliquots of the suspension were treated with 10% chloroform, vortexed for 1.5 minutes, then centrifuged for 1 minute at 14,000rpm. The supernatant containing the phage was aliquoted into clean 2mL Eppendorf tubes, and the process repeated a second time to ensure the complete removal of bacterial hosts.

The number of infectious phage present was quantified using plaque assays. ISP was 10-fold serially diluted in 1X minimal media (M9) from 10^0^ to 10^−8^. 100μL of overnight culture of 13S44S9 and 50μL of diluted phage were added to 5mL of soft LB agar, gently mixed, and poured over a 20mL LB agar plate. Plates were left to dry before being inverted and incubated at 37°C for 24 hours. Plaques were counted and plaque forming units (PFU)/mL determined based on the dilution with the highest number of discernible plaques. Each plaque assay was repeated at least 3 times to account for plating error.

### Assessing susceptibility using plaque assays

10-fold serial dilutions of ISP were prepared in 1X M9, ranging from 10^0^ to 10^−8^. Bacterial isolates were diluted in LB, plated out to a final density of 2×10^10^ bacteria/mL, and left to dry before 5μL of each ISP dilution was spotted on each LB agar plate. The plates were then inverted and stored at 37°C. Plaques were counted after 24-hours and converted to PFU/μL, based on the dilution with the highest number of discernible plaques. Of the 64 bacterial hosts tested, 6 biological replicates were obtained for 42 strains, 5 replicates for 12 strains, and 4 replicates for 10 strains.

### Assessing susceptibility using optical density assays

In each well of a 96-well plate, 180μL of LB broth was added alongside 10μL of each *Staphylococcus* isolate at a concentration of 1×10^6^ CFU/mL. For infected plates, 10μL of ISP was then added to each well at an MOI of 0.01, whereas for control plates 10μL of 1X M9 was added. The layout of the plates was randomised for each block to minimise the influence of position effects but remained the same between the infected and non-infected plates within a block. Three LB media and three 1X M9 controls were added to each plate to check for contamination. The plates were sealed with an adhesive PCR plate seal (Thermo Scientific) and incubated for 24-hours at 37°C, 180rpm. Following the incubation, OD was read at 600nm on a MultiSkan Sky Microplate Spectrophotometer (Thermo Fisher). To standardise for any differences in bacterial growth rate, susceptibility was calculated as a proportion change in OD due to infection, as follows:

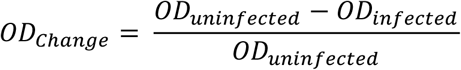

Several collected data points reported a proportion change in OD less than zero. As this was likely the result of anomalous OD readings, outlier analysis was performed. Datapoints more than 1.5 times the interquartile range outside the upper and lower quartile of the data were considered minor outliers, and those more than 3 times the interquartile range outside the upper and lower quartiles were considered major outliers. All analyses included here were robust to the removal of both major and minor outliers (Table S2-5), and so only data with all major and minor outliers removed is presented outside of the supplementary data. Following outlier removal, 6 biological replicates were obtained for 56 of the bacterial strains tested, 5 replicates were obtained for 7 of the strains, and 3 replicates were obtained for 1 strain.

### Assessing susceptibility using quantitative PCR

Infections used to measure susceptibility using quantitative (q)PCR were set up identically to those in the OD assays described above. Following the 24-hour incubation, 100μL of each sample was transferred to a screw cap microcentrifuge tube and heat treated at 90°C for 10-minutes to inactivate the bacteria. Additionally, 12 samples of the aliquot of ISP used to infect the *Staphylococcus* strains were extracted to provide an initial viral load at infection timepoint zero. To extract the viral DNA, 100μL of 10% w/v Chelex 100 (Merck Life Sciences, UK), 2ul of 20ng/μL Proteinase K (Merck Life Sciences, UK), and ∼10 1mm zirconia beads (Thistle Scientific, UK) were added to each sample before mechanically lysing the bacteria using an Omni Bead Ruptor 24 (Camlab, UK). Samples were spun down and heat treated for 10 minutes at 95°C to inactivate the Proteinase K before being spun down for a further 5 minutes to sediment out the Chelex. For each sample, 50μL of the supernatant containing bacterial and viral DNA was transferred and stored at -20°C.

qPCR was performed on each of the samples using an Applied Biosystems StepOnePlus system with a Sensifast Hi-Rox Sybr kit (Bioline). Cycle conditions were as follows: initial denaturation at 95°C for 120 seconds, then 40 cycles of 95°C for 5 seconds, and 60°C for 30 seconds. ISP was measured using the following primers: forward, 5’-CCTGTACCGGCTTGACTCTC -3’; reverse, 5’-AGCTACAACCGAGCAGTTAGA -3’, which were confirmed to have a near 100% efficiency when tested in the presence of each bacterial isolate. Pilot experiments showed that normalisation to either *Staphylococcal* genomic DNA or an exogenous DNA spike made little difference to the between sample variation in ISP viral load and was therefore not used.

For each sample, two technical replicates of the qPCR reaction were performed. Amplification of the correct product was confirmed by melt curve analysis: samples that had failed to amplify the product, showed evidence of melt curve contaminants, or departed from the melt curve peak of positive samples by ±1.5°C were excluded. Correction for between-plate effects between technical replicates was performed using a linear model as previously described (79,80). Mean viral C_t_ values from technical replicate pairs (C_t:24_) were normalised to an initial dose of ISP (C_t:0_) and converted to fold change in viral load using the 2^−ΔCt^ method, where ΔCt = C_t:24_ – C_t:0_. These values were then converted to a log_10_ fold change in viral load over 24-hours prior to analysis. Of the 64 bacterial strains tested, 6 biological replicates were obtained for 58 strains, 5 replicates for 5 strains, and 4 replicates for 1 strain.

### Inferring the host phylogeny

To infer the evolutionary relationships between *Staphylococcus* isolates, a core genome phylogeny was constructed using BEAST v1.10 (81). Briefly (see Supplementary Methods for full details), whole genome sequences were collected for 8 previously sequenced strains from NCBI (Supplementary Table 1), and the remaining 56 sequences were obtained through whole genome sequencing. Library preparations and sequencing were performed by MicrobesNG (Birmingham, UK). Genomic DNA libraries were prepared using the Nextera XT Library Prep Kit (Illumina, San Diego, USA) with twice the stated amount of input DNA and PCR elongation increased to 45-seconds. Pooled libraries were quantified using the Kapa Biosystems Library Quantification Kit for Illumina and sequenced using Illumina sequencers (HiSeq/NovaSeq) with a 250-bp paired end protocol. Sequence reads were deposited on NCBI under the BioProject ID: PRJNA894984 (see Supplementary Table 1). Genomes were assembled and quality controlled as described in the Supplementary Methods.

D*e novo* assemblies were annotated using Prokka (v2.8.2) (82) and orthologous genes identified with Panaroo (83) using a sequence identity threshold of 0.7. Orthologous genes were then used to generate a core genome alignment of the 102 genes shared between each of the 64 *Staphylococcus* isolates. Phylogenetic trees were constructed using BEAST v1.10 (81). Sequence alignments were fitted to a HKY substitution model using relaxed uncorrelated molecular clock models, gamma distributions of rate variation, and constant population size coalescent priors (81). Separate substitution models and molecular clocks were fitted to 1^st^/2^nd^ and 3^rd^ codon positions, to reflect differences in selective constraint (84). Two independent MCMC chains were run for each model until both convergence and a <10% burn-in was achieved. Convergence of all parameters was checked using Tracer v1.6 (85).

The within-*S. aureus* and among-species phylogenies were constructed by dropping the relevant tips from the 64-strain tree using the R package ape (86). To ensure that the phylogenetic relationship between species and strains were accurately resolved in the 64-strain tree, separate phylogenetic trees were constructed using only the *S. aureus* samples or species isolates with 13S44S9 as a representative *S. aureus* strain (see Supplementary Methods). As the topology of the smaller phylogenies matched that of the phylogenies made by dropping tips from the whole phylogeny, the dropped tip phylogenies were used for our analyses as they had comparable branch lengths to the 64-strain phylogeny.

### Phylogenetic mixed models

Phylogenetic generalised linear mixed models were used to investigate the effect of host relatedness on susceptibility to infection with ISP, and to examine correlations between the methods used to assess susceptibility. Multivariate models were fitted using the R package MCMCglmm (87) with susceptibility, as determined by each method (plaque assay, OD, and qPCR), as the response variable. Unscaled trees were used to correct for differences in the evolutionary divergence of our within- and among-species trees.

The structures of the model were as follows:

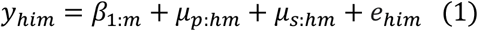

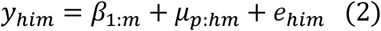

In these models, *y*_*him*_ is the susceptibility measured by method *m* in the *i*^*th*^ biological replicate of host *h*. The fixed effect *β*_1_ represents the intercepts for each method, the random effect *μ*_p_ represents the effects of the host phylogeny assuming a Brownian model of evolution, and *e* represents the model residuals. Model (1) also includes a strain-specific random effect that is independent of the host phylogeny (*μ*_s:*hm*_). This explicitly estimates the non-phylogenetic component of between-strain variance and allows the proportion of variance explained by the host phylogeny to be calculated. *μ*_s:*hm*_ was removed from model (2) as model (1) failed to separate the phylogenetic and strain-specific effects. The effect of evolutionary distance of each host from the amplification host (*S. aureus* 13S44S9) was added to the model as a fixed effect (*β*_2:hm_). This was done to examine if the evolutionary distance from the amplification host explained any observed differences in susceptibility between strains.

Susceptibilities measured by optical density (OD) and qPCR were treated as normally distributed. However, to account for the zero-inflated nature of the plaque assay data, plaque assay susceptibility was divided into two separate variables – equivalent to a hurdle modelling approach – with a binary variable indicating whether the bacteria was permissive (1) or non-permissive (0) to infection, modelled using probit link, and conditional on being permissive, a continuous variable containing PFU/μL which was treated as normally distributed.

Within each of these models, the random effects and residuals were assumed to follow a multivariate normal distribution with a mean of 0 and covariance structure **V**_*p*_⊗**A** for the phylogenetic effects, **V**_*s*_⊗**I** for strain-specific effects, and **V**_*e*_⊗**I** for residuals, where ⊗ represents the Kronecker product. **A** represents the host phylogenetic relatedness matrix, **I** an identity matrix, and **V** represents 4 × 4 covariance matrices describing the variances and covariances in susceptibility for the different methods. Specifically, the matrices **V**_*p*_ and **V**_*s*_ describe the phylogenetic and non-phylogenetic between-strain variances in susceptibility for each method and the covariances between them, whereas the residual covariance matrix **V**_*e*_ describes the within-strain variance that includes both true within-strain effects and measurement errors. Because each biological replicate consists of a measurement from a single method, the covariances of **V**_*e*_ cannot be estimated and were set to 0. Additionally, the residual variance for the binary variable cannot be estimated and was fixed at one. An additional version of model (1) was run with evolutionary distance from the amplification host added as a fixed effect (*β*_*d:hm*_), but this was shown to be non-significant.

Models were run for 13 million MCMC generations, sampled every 5,000 iterations with a burn-in of 3 million generations. Parameter expanded priors were placed on the covariance matrices, resulting in multivariate F distributions with marginal variances being scaled by 1000. Inverse-gamma priors were placed on the residual variances, with a shape and scale equal to 0.002. To ensure the model outputs were robust to changes in prior distribution, models were also fitted with inverse-Wishart priors, which gave qualitatively similar results.

The proportion of between strain variance that can be explained by the phylogeny was calculated from model (1) using the equation 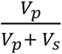, where *V*_*p*_ and *V*_*s*_ represent the phylogenetic and strain-specific components of between-strain variance respectively, and is equivalent to phylogenetic heritability or Pagel’s lambda (88,89). Phylogenetic heritability was additionally calculated from model (1) as the proportion of the total variance explained by phylogeny, using the equation 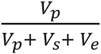, where *V*_*e*_ represents the within-strain variance. The repeatability of susceptibility measurements was calculated from model (2) as 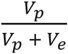 where *V*_*e*_ is the residual variance (87). Inter-strain correlations in viral load between each method were calculated from model (2) *V*_*p*_ matrix as 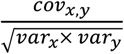 and the slopes (*β*) of each relationship as 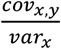. Parameter estimates stated below are means of the posterior density, and 95% credible intervals (CIs) were taken to be the 95% highest posterior density intervals.

### Leave-one-out cross-validation

To investigate the ability of the bacterial host phylogeny to predict the susceptibility of an novel host, leave-one-out cross-validation was used (90), whereby multiple versions of model (1) were fitted, each with the data from a single bacterial strain removed, and the model challenged to predict the susceptibility of the “unknown” host given only its evolutionary relationships to other *Staphylococcus* strains and their measured susceptibilities. Additionally, a null model was run, whereby the phylogeny and all information on the relationship between host species was removed from the leave-one-out cross-validation model. Prediction errors from the leave-one-out cross-validation and the null model were compared using Wilcoxon rank sum tests (91) to determine whether information on the relationship between host species significantly improved the ability of the model to predict the susceptibility of an unknown host.

### Data availability

All data files and R scripts used in this study are available in an online repository: https://doi.org/10.6084/m9.figshare.21642209.v1.

## Results

### ISP is a broad host range phage with varying infectivity across *Staphylococcus* species

To investigate the ability of the host phylogeny to explain variation in susceptibility to virus infection in bacteria *–* and how different methods may vary in their quantification of phage host range – we experimentally infected 64 strains of *Staphylococcus* with the bacteriophage ISP and assessed susceptibility using three distinct methods: plaque assays, OD assays, and qPCR. When assessed using plaque assays, 64% of host strains were seen to be permissive to infection with ISP. However, both OD and qPCR measures of susceptibility showed that ISP was capable of infecting 97% of the host panel (Figure 1). Variation in susceptibility between *Staphylococcus* isolates was seen in every method: the mean PFU/μL of permissive hosts ranged from 2.2×10^3^ in *S. simulans* to 8.1×10^5^ in *S. aureus* strain JW31330OBHY1; the mean proportional decrease in OD ranged from 0.01 in *S. aureus* strain SAR1218N1 to 0.90 in *S. aureus* strain B142S1; and the mean change in qPCR viral load ranged from a 1.2-fold increase in *S. simiae* to a ∼300,000-fold increase in *S. aureus* strain DAR06181LC1 (Figure 1). Together, these results show that ISP is both able to infect a broad range of *Staphylococcus* species, and that susceptibility to ISP varies widely across the *Staphylococcus* genus.

**Figure 1:**
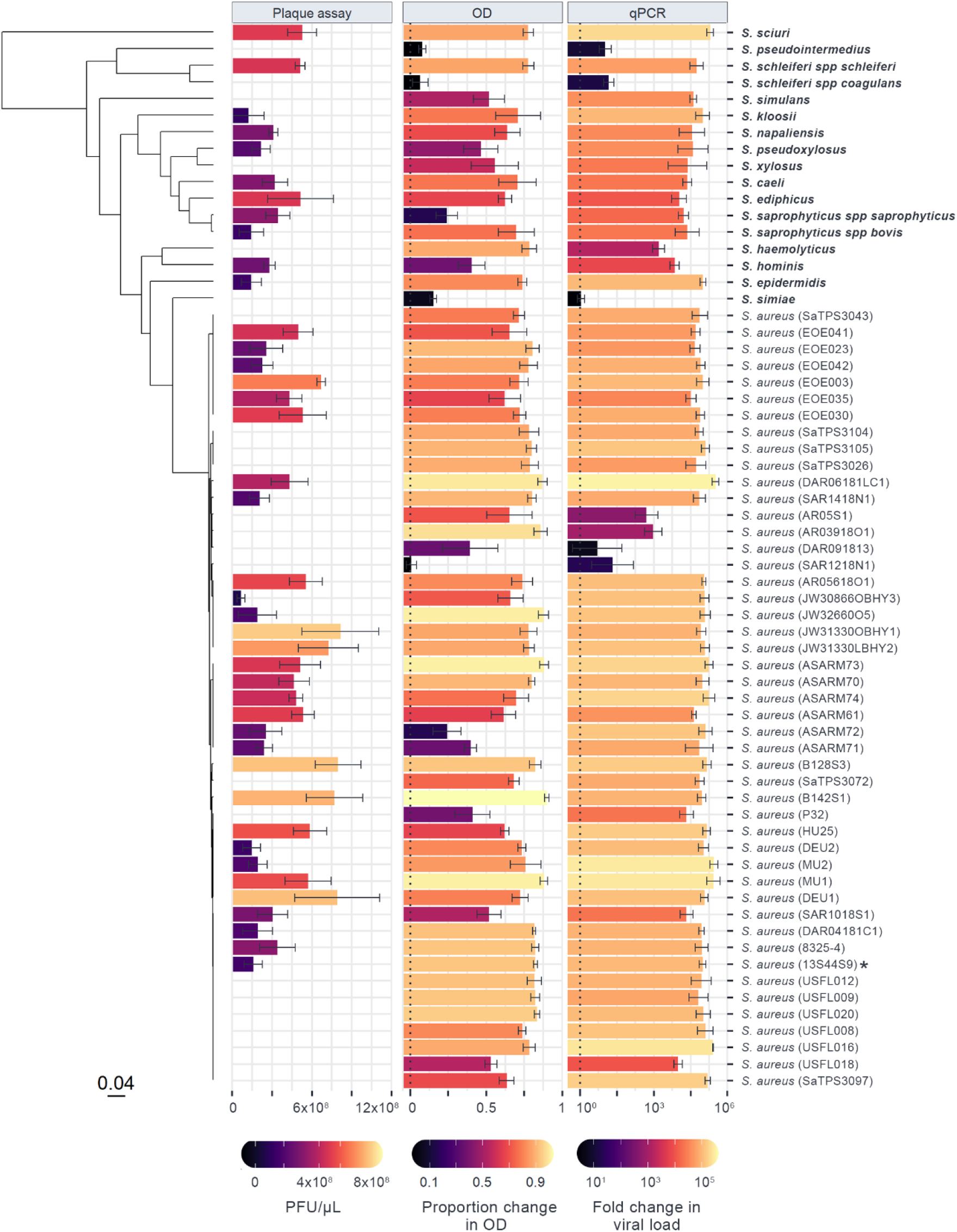
Susceptibility to ISP across a panel of 64 *Staphylococcus* isolates, measured using three distinct methods. Bar lengths and colour show the mean change in ISP assessed by plaque assay (PFU/μL), optical density (proportion decrease in OD with infection after 24 hours), and qPCR (log_10_ fold change in viral load after 24 hours), with error bars representing the standard error of the mean across at least four biological replicates. The phylogeny of *Staphylococcus* hosts is presented on the left, with the scale bar representing the number of nucleotide substitutions per site. Strain names are presented on the right, with non-aureus species in bold and the amplification host labelled with an asterisk (a full version of the tree is available at https://doi.org/10.6084/m9.figshare.21642209.v1).

### Susceptibility to ISP across *Staphylococcus* is explained by the host phylogeny

The phylogeny of *Staphylococcus* host species inferred here is broadly consistent with previous studies of these taxa (84), with the close phylogenetic relationships between species being generally well supported and hosts falling into two main clades (Figure 1). The *Staphylococcus* phylogeny is characterised by high susceptibility to ISP across a large number of isolates, with smaller clades showing reduced susceptibility to infection (i.e., the clade containing *S. aureus* strains AR05S1, AR03918O1, and DAR091813). Some clades show intermediate levels of susceptibility (e.g., the clade containing *S. haemolyticus* and S. *hominis*) while some contain both permissive and non-permissive strains (e.g., the clade containing *S. aureus* strains SAR1218N1 and AR05618O1).

Phylogenetic generalised linear mixed models were fitted to the data to determine the proportion of variation in susceptibility explained by the host phylogeny. Estimates of repeatability and phylogenetic heritability for the binary plaque assay, OD, and qPCR data were close to 1 with narrow credible intervals. The convergence of phylogenetic heritability and repeatability estimates for the binary plaque assay, OD, and qPCR data at 1 suggests that the between-strain phylogenetic component explains a high proportion of the variation in susceptibility with little within-strain variation or measurement error (Table 1). Estimates of repeatability and phylogenetic heritability for the continuous (PFU/μL) component of the plaque assay were poor, with wide credible intervals spanning 0-1. The effect of phylogeny on the susceptibility of *Staphylococcus* hosts to ISP can be further seen in ancestral state reconstructions (Figures 2 and 3), where different clades are seen to have similar susceptibilities to infection. This is particularly apparent within-*S. aureus* samples, where clades show either high, intermediate, or low susceptibility to ISP (Figure 3).

**Table 1:** Estimates for the repeatability and phylogenetic heritability across 64 *Staphylococcus* isolates. Estimates of repeatability are taken from model (2) and estimates of phylogenetic heritability (the variation explained by the host phylogeny) are taken from model (1). * indicates the phylogenetic heritability calculated as the proportion of variation that is attributed to phylogeny divided by the total variation (phylogenetic, non-phylogenetic between-strain, and within-strain variation) as opposed to phylogenetic and non-phylogenetic between-strain variation. PA = plaque assay, CI = credible interval.

| Method | Repeatability |  | Phylogenetic heritability |  | Phylogenetic heritability of total variance* |  |
| --- | --- | --- | --- | --- | --- | --- |
|  | Mean | 95% CI | Mean | 95% CI | Mean | 95% CI |
| Binary PA | 1.00 | 1.00, 1.00 | 1.00 | 1.00, 1.00 | 1.00 | 1.00, 1.00 |
| Continuous PA | 0.00 | 0.00, 0.00 | 0.68 | 0.01, 1.00 | $6.54 \times 10^{-6}$ | $1.91 \times 10^{-15}$, $1.43 \times 10^{-5}$ |
| OD | 0.98 | 0.96, 0.99 | 0.97 | 0.94, 1.00 | 0.94 | 0.88, 0.98 |
| qPCR | 0.99 | 0.99, 1.00 | 1.00 | 1.00, 1.00 | 0.99 | 0.98, 1.00 |

**Figure 2:**
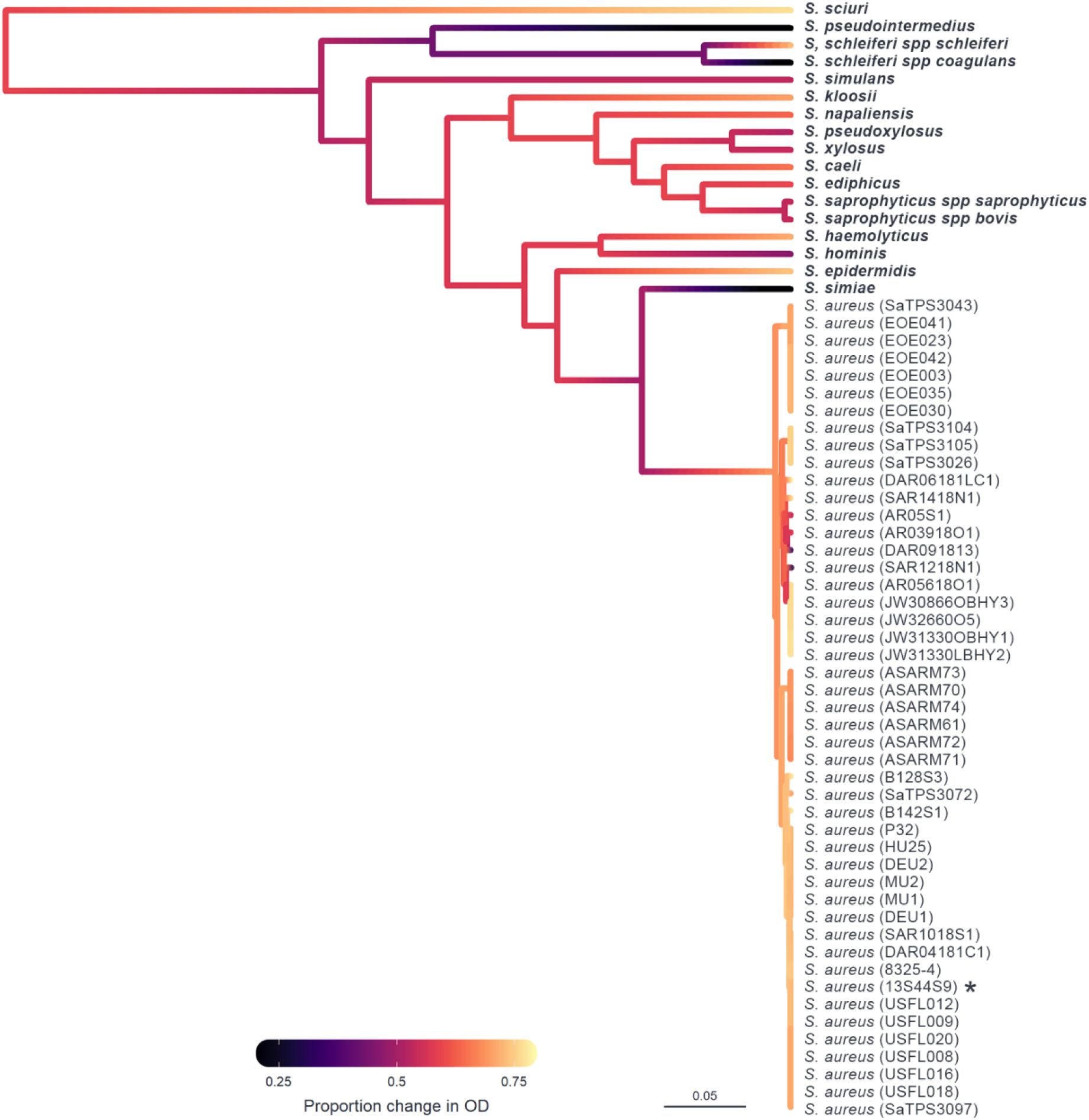
Ancestral state reconstruction of susceptibility to ISP measured by OD. Ancestral states were estimated from model (2) for each node and plotted in colour across the *Staphylococcus* host phylogeny, with the scale bar representing nucleotide substitutions per site. Colours represent susceptibility of a host to infection with ISP measured by OD (proportion change in optical density in infected compared to non-infected cultures), with black representing the lowest level of susceptibility and yellow the highest. Strain IDs are presented on the right, with non-aureus species in bold and the amplification host labelled with an asterisk (*).

**Figure 3:**
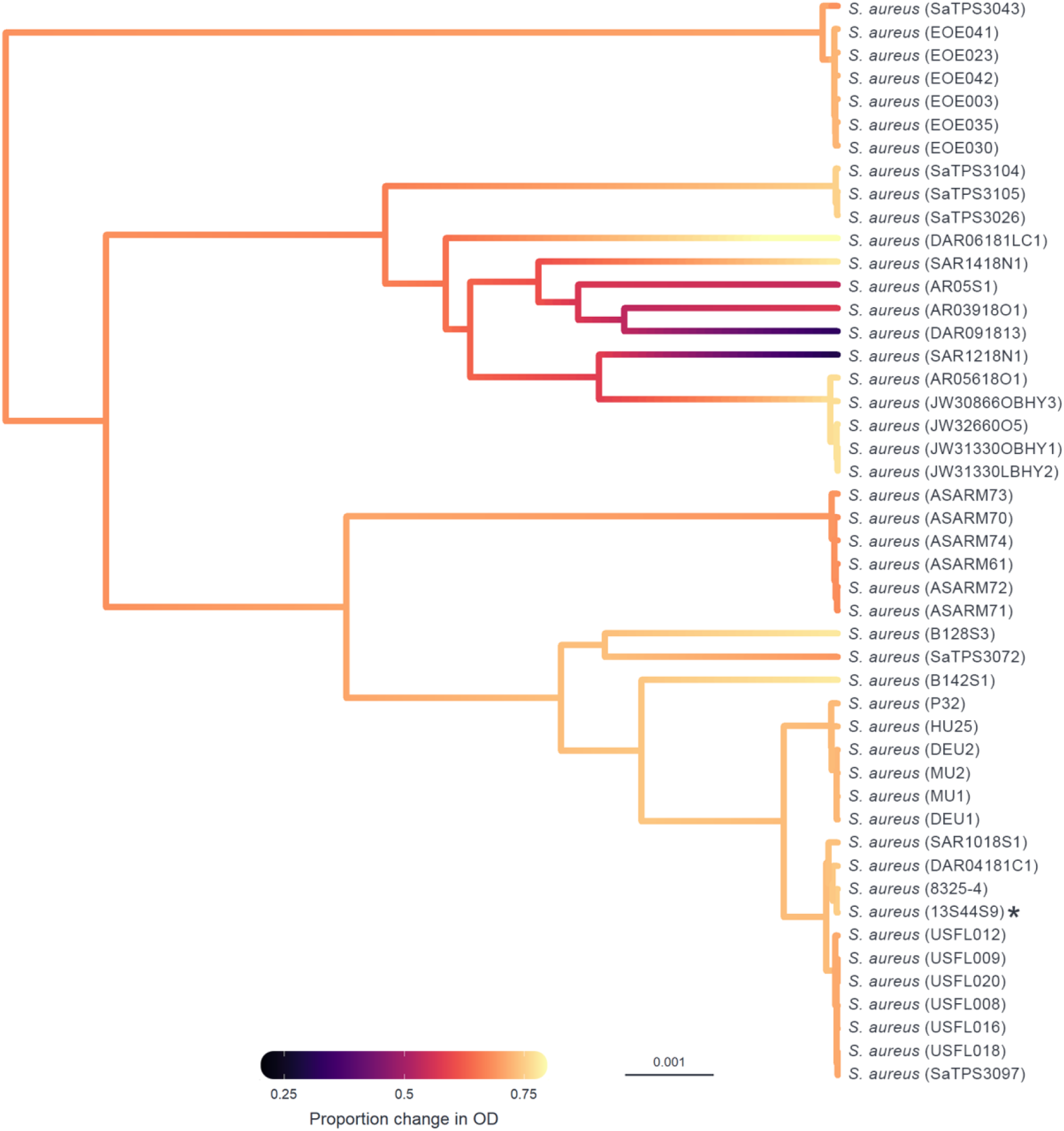
Ancestral state reconstruction of the *S. aureus* host strains susceptibility to ISP measured by OD. Ancestral states were estimated from model (2) for each node and plotted in colour across the *Staphylococcus aureus* host phylogeny, with the scale bar representing nucleotide substitutions per site. Colours represent susceptibility of a host to infection with ISP measured by OD (proportion change in optical density in infected compared to non-infected cultures), with black representing the lowest level of susceptibility and yellow the highest. Strain IDs are presented on the right and the amplification host labelled with an asterisk (*).

To determine if the observed phylogenetic signal is consistent across evolutionary scales, we reduced the phylogeny to one containing only the *S. aureus* samples (Table S6) and one containing each of the *Staphylococcus* species (Table S7). Similar estimates of repeatability were observed for both the within-*aureus* and among-species phylogeny models. However, both models showed a reduced ability to estimate phylogenetic effect, with wide credible intervals around estimates (apart from estimate for qPCR in the within-*aureus* model). It is likely that the observed difference in ability to estimate heritability between the whole phylogeny model compared to the reduced phylogeny models is down to reduced statistical power, causing the latter models to struggle to separate the phylogenetic and strain-specific effects.

Finally, to ensure that ISP had not adapted to its amplification host prior to the experiment, we looked for an effect of distance from the amplification host (*S. aureus* strain, 13S44S9). No effect of distance from the amplification host on the susceptibility of *Staphylococcus* to ISP was found (*β* = 0.03, 95% credible interval: -1.46, 1.56), suggesting that the observed phylogenetic signal was not being driven by adaptation of ISP to the amplification host.

### Measures of susceptibility from plaque assays, OD assays, and qPCR are positively correlated across hosts

Correlations between plaque assay, OD, and qPCR measures of susceptibility to ISP across bacterial host strains were estimated from the variance-covariance matrices of model (2) (Table 2). A strong positive inter-strain correlation was observed between susceptibility measured by OD and qPCR (*r* = 0.97, 95% CI: 0.92, 1:00), and a strong positive correlation was observed between the binary measure of plaque assay and OD (*r* = 0.94, 95% CI: 0.86, 1.00) and the binary measure of plaque assay and qPCR (*r* = 0.98, 95% CI: 0.94, 1.00). However, no evidence of a correlation was observed between the continuous plaque assay data and either OD or qPCR, with correlation coefficients approximately zero and credible intervals spanning -1 to 1 (Table 2). Similar correlations were observed between methods for the within *S. aureus* (Table S8) and among-species (Table S9) phylogenies, with the qPCR:OD correlation being the only one consistently and significantly positive. A correlation between the binary measure of PA and the other measures of susceptibility was observed in the among-species phylogeny but not the within-*S. aureus* phylogeny, suggesting that the correlation observed between these measures is being driven by strong correlations seen at the species level rather than the within-species level.

**Table 2:** Inter-strain correlations in susceptibility measures between methods. Numbers show the mean estimates for the correlation strength (*r*, white cells) and slope (*β*, grey cells) between pairs of methods, with 95% credible intervals (CIs) indicated in brackets. The correlations and slopes were calculated with the variables in columns as *x* and variables in rows as *y*. Estimates with CIs that do not span zero are highlighted in bold. PA = plaque assay, *value on a probit scale.

|  | Binary PA | Continuous PA | OD | qPCR |
| --- | --- | --- | --- | --- |
| Binary PA | - | - | **0.94**<br>**(0.86, 1.00)** | **0.98**<br>**(0.94, 1.00)** |
| Continuous PA | - | - | -0.01<br>(-0.99, 0.99) | -0.01<br>(-1.00, 1.00) |
| OD | **0.50\***<br>**(0.50, 0.50)** | -0.00<br>(-0.12, 0.17) | - | **0.97**<br>**(0.92, 1.00)** |
| qPCR | **0.51\***<br>**(0.50, 0.52)** | 0.01<br>(-1.30, 0.97) | **7.45**<br>**(5.53, 9.59)** | - |

### Mixed evidence for host phylogeny improving the accuracy of susceptibility predictions

As the host phylogeny explains a large proportion of the variation in ISP susceptibility, it may allow for the susceptibility of untested *Staphylococcus* strains and species to be predicted based on their evolutionary relationships to *Staphylococci* with known susceptibilities. To investigate the ability of the bacterial host phylogeny to predict susceptibility, leave-one-out cross-validation was used, whereby multiple versions of model (1) were fitted, each with the data from a single bacterial strain removed, and the model challenged to predict the susceptibility of the “unknown” host given only its evolutionary relationships to other *Staphylococci* strains and their measured susceptibilities (Figure 4).

**Figure 4:**
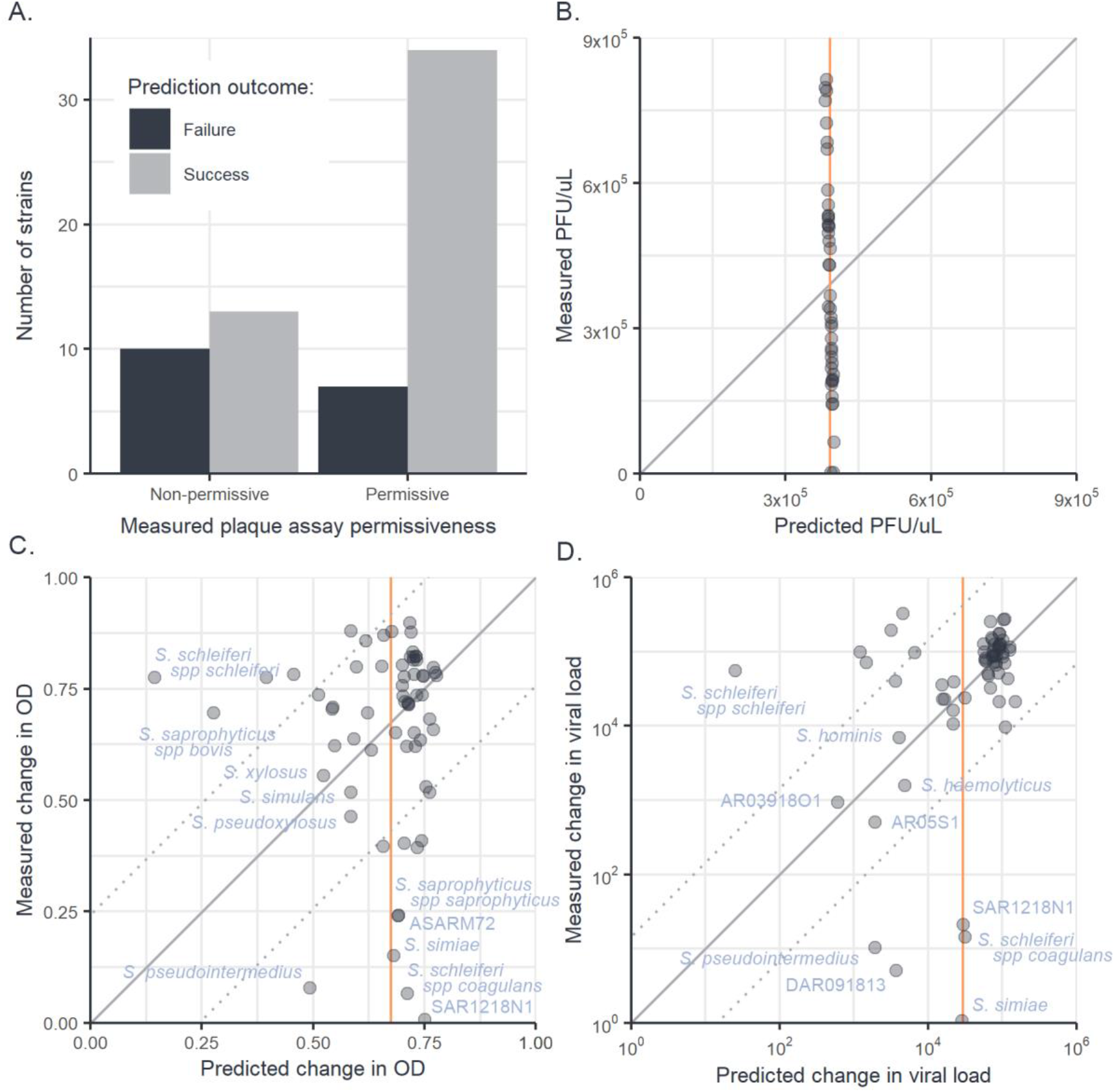
Leave-one-out cross-validation of phylogenetic mixed models fitted to plaque assay, OD, and qPCR data. Predictions of “unknown” trait values from a leave-one-out cross-validation of phylogenetic model (2) for binary plaque assay (A), continuous (PFU/μL) plaque assay (B), change in OD with infection (C) and fold changes in viral load measured by qPCR (D). Each datapoint represents an individual strain of *Staphylococcus* whose measured trait value has been removed from model (2) and predicted from its evolutionary relationships to other *Staphylococcus* isolates and their trait values. Solid diagonal lines illustrate the location of 1:1 predictions and dotted lines indicate the root-mean-squared errors around these lines. Orange vertical lines represent the predicted trait values of all strains from a null (intercept only) model.

For the binary component of plaque assay, the phylogenetic model was able to correctly predict a strain’s permissiveness to ISP infection in 73% of cases (Figure 4A), whereas a null model without phylogeny predicted all hosts to be permissive, achieving an accuracy of 63%. When asked to predict the continuous (PFU/μL) component of plaque assays, errors from the phylogenetic and null models were indistinguishable (Wilcoxon rank sum test: W_43,43_ = 886, p = 0.74), as in the phylogenetic model most of the variation in PFU/μL was partitioned into the model residuals, causing trait values predicted from the phylogenetic component to cluster around the across-strain mean (Figure 4B). When asked to predict susceptibility measured by OD assay (Figure 4C), the errors produced by the phylogenetic model were also not significantly different from those produced by a null model (Wilcoxon rank sum test: W_64,64_ = 2111, p = 0.77). However, when predicting viral loads from qPCR data (Figure 4D), the phylogenetic model produced a small but significant decrease in error compared to the null model (Wilcoxon rank sum test: W_64,64_ = 2726, p < 0.01).

In the OD and qPCR assays most strains were both similar in susceptibility to their close relatives and showed susceptibility close to the across-strain mean (Figure 1, 4C-D), causing the phylogenetic and null models to produce similar predictions for the majority of strains. In cases where hosts existed in clades with susceptibilities different from the across-strain mean, prediction accuracy increased when phylogeny was included (e.g., qPCR prediction for *S. haemolyticus & S. hominis*, OD prediction for *S. xylosus* and *S. pseudoxylosus*). However, when hosts had susceptibility near the across-strain mean and had close relatives that differed from this mean, the phylogenetic model produced larger errors in prediction than the null model (e.g., both OD and qPCR predictions for *S. schleiferi spp schleiferi*). Lastly, where hosts differed from the across-strain mean but had close relatives that conformed to the mean, both phylogenetic and null models were poor predictors of susceptibility (e.g., both OD and qPCR predictions for *S. schleiferi spp coagulans* and *S. aureus* SAR1218N1). Together, these results suggest that the host phylogeny may allow for the limited prediction of host susceptibility in some cases, but may also mislead predictions in host strains that differ strongly from the susceptibilities of their close relatives.

## Discussion

Closely related host species present similar environments to novel viruses (88,92), and so tend to share similar levels of susceptibility (17,19–21,26). Here, we have examined how susceptibility to a broad host range phage varies across a diverse panel of *Staphylococcus* bacteria and determined what proportion of the variation in phage susceptibility is explained by the relationships between bacterial hosts. ISP was capable of infecting 97% of the host strains investigated here when measured by OD and qPCR assays, but only 64% of these strains appeared permissive when tested by plaque assay. We found that variation in susceptibility across our S*taphylococcus* panel – measured using OD, qPCR, and the binary component of plaque assays – was largely explained by the host phylogeny, and these effects were seen at both within-species, and among-species phylogenetic scales. No effects of evolutionary distance from the amplification host were seen, with the patterns seen instead being driven by the existence of host clades sharing similar levels of susceptibility to ISP. Strong positive correlations were seen between the susceptibility measures from the binary PA, OD, and qPCR assays, but not between these methods and the PFU/μL measures taken from plaque assays. Together, these results suggest that ISP has a broad ability to infect bacterial hosts within the *Staphylococcus* genus, and that a large proportion of the variation in susceptibility to ISP across *Staphylococcus* hosts can be explained by the evolutionary relationship between bacterial hosts.

Historically, phages were believed to have narrow host ranges consisting of one, or a few, closely related host species. However, recent studies have shown through metagenomic sequencing and environmental sampling that broad host range phage are more common than previously thought (93). Here we showed that ISP has a broad host range, able to infect both within and among *Staphylococcus* species. Many studies have focused on the molecular mechanisms of generalism at the attachment stage, for example polyvalent phage (that can bind to multiple host receptors) (94–96) or phage that bind a single, conserved receptor are more likely to have a broad host range (97). However, generalism may also be dependent on the ability of a phage to subvert the phage resistance mechanisms of multiple host species (93). Permissive *Staphylococcus* strains showed wide variation in their susceptibility to ISP, with some strains showing small decreases in OD and low viral load, and others showing large decreases in OD and high viral loads. This among-strain and among-species variation in susceptibility may be due to differences in the affinity and availability of receptors for ISP adsorption on the bacterial cell surface, or variation in the efficacy of bacteria host immune defences (98). Identification of the ISP receptor, as well as the host pathways involved in generating variation in susceptibility to ISP across bacteria, are important next steps in building a more complete understanding of ISP infection and how it may be implemented in therapies against multiple bacterial pathogens.

Bacterial evolution is known to involve several mechanisms that could disrupt the phylogenetic heritability of phage immunity. Notably, cell surface receptors can be easily altered by point mutations and components of phage resistance may be frequently gained and lost through horizontal gene transfer, which can lead to high levels of variation in the phage defence systems present in bacterial genomes (33,99–103). Despite the potential for horizontal gene transfer and large effect point mutations to disrupt the phylogenetic signal in phage susceptibility, our models suggest that the evolutionary relationships between bacterial hosts can capture a large proportion of the variation in susceptibility to ISP. If horizontal gene transfer is occurring more frequently between closely related strains of *Staphylococcus*, then their influence on phage susceptibility may be phylogenetically conserved and this variation may have been captured in the core genome phylogeny. In any case, our results indicate that more closely related *Staphylococcus* strains and species are more likely to share similar susceptibilities to phage infection, consistent with patterns seen in previous studies of animal hosts and viruses (17,19–21,26). These patterns may be the result of different host clades having acquired or lost immune or cellular components that affect susceptibility to infection in a common ancestor, although we lack the ability to identify these specific components here.

In this study, we used three methods to assess the susceptibility of *Staphylococcus* to a bacteriophage: plaque assays, optical density assays, and qPCR. Susceptibility measured by binary PA, OD, and qPCR were strongly positively correlated. However, no correlation was seen between the continuous component of plaque assay and either other method. While OD and qPCR suggest a broad host range for ISP, plaque assays underestimated the susceptibility of the host panel, with only 63% of strains appearing permissive to infection. Historically, plaque assays were thought to overestimate bacteriophage host range due to large numbers of phage adsorbing to and lysing the bacterial cell or the effects of residual endolysins or bacteriocins in the phage stock (104). This discrepancy may be due to variation in the ability of ISP to induce lysis in solid media compared with liquid culture. However, as OD and qPCR show a strong correlation in their measurement of susceptibility and their host range estimations are similar, it is likely they are providing a more accurate estimation of host range. The strong correlation between OD and qPCR may be expected as they measure infection under similar conditions, OD and qPCR assess infection while bacteria are planktonic whereas PA assess susceptibility under static conditions. *Staphylococcus* bacteria have been shown to have different gene expression when planktonic compared with when they form biofilms, including the expression of virulence factors (105–108), which may explaining some of the observed variation in susceptibility between the different assays. Interestingly, we observe a strong positive correlation between OD and qPCR even though both assays are measuring different aspects of infection, with OD providing a measure of damage done to the host and qPCR a measure of virus replication and persistence. That these two infection traits are strongly correlated suggests that *Staphylococcus* phage immunity is strongly tied to its ability to disrupt and prevent phage replication, as opposed to mitigating damage while tolerating phage replication.

Being able to predict and prevent emerging infectious diseases is a major goal of scientific research, and one of the first steps of that process is being able to predict whether a host will be susceptible to a novel pathogen (109). While our phylogenetic models suggest that a high proportion of the variation in susceptibility between host species can be attributed to the relationship between hosts, they showed a limited ability to predict the susceptibility of a host given only its relationship to other hosts and their susceptibilities. Several additions may improve the accuracy of predictions of susceptibility using these models. Firstly, improving the depth of our phylogeny – i.e., adding more strains that reduce the number of isolated species on long evolutionary branches – would increase the number of observations that the model can use to predict susceptibility. For example, *S. simiae* is isolated within the phylogeny, with few close relatives, and had a poorly estimated susceptibility due to the distance between it and its nearest neighbour. Secondly, furthering our understanding of the mechanisms by which susceptibility can change in a non-phylogenetically tractable way may improve our ability to predict susceptibility when phylogeny is not informative. Finally, alternative approaches to the prediction of susceptibility and other related infection traits using genome composition bias have been shown to consistently outperform phylogenetic mixed models (109–111), suggesting that there are further metrics that could be incorporated into models attempting to predict susceptibility.

Overall, our results demonstrate that the bacterial host phylogeny is an important determinant of phage susceptibility and replication across novel hosts, and that the relationship between host species may be a useful predictor of virus host range both in the context of emerging infectious disease and bacteriophage therapy. Further work is required to understand the specific interactions underlying variation in ISP susceptibility across bacterial hosts; the relationship between host damage, virus replication, and virus persistence in this system; and how the patterns of phage susceptibility across bacterial phylogenies may vary under different infection conditions and contexts.

## Acknowledgements

Staphylococcus isolates were kindly provided by Edward Feil and Kay Fountain from the University of Bath and Gavin Paterson at the University of Edinburgh. ISP was originally isolated by the Eliava Institute (Tbilisi, Georgia) and kindly provided by Jean-Paul Pirnay and Maya Merabishvili at the Queen Astrid Military Hospital (Brussels, Belgium). Genome sequencing was provided by MicrobesNG (http://www.microbesng.com). S.K.W. is supported by a studentship funded by the Biotechnology and Biological Sciences Research Council (BBSRC) South West Biosciences Doctoral Training Partnership (BB/M009122/1). M.M. was funded by a PhD from the Medical Research Council with support from the Raymond and Beverly Sackler Fund. L.A.W. is funded by a Sir Henry Dale Fellowship through the Royal Society and Wellcome Trust (109385/Z/15/Z). J.D.H. is supported by NERC (NE/P000924/1). A.B. is supported by NERC (NE/S000771/1 and NE/V012347/1). B.L. and R.M.I are supported by a Sir Henry Dale Fellowship jointly funded by the Wellcome Trust and the Royal Society (109356/Z/15/Z).

For the purpose of Open Access, the author has applied a CC BY 4.0 public copyright licence to any Author Accepted Manuscript version arising from this submission.

## Author contributions

Provided resources: GKP.

Conceived and designed experiments: SKW, RMI, BL, AB.

Performed the experiments: SKW.

Genome assembly and phylogenetic analysis: SKW, MM, LAW.

Statistical analysis: SKW, RMI, JDH, BL.

Wrote the paper: SKW, RMI, BL, AB with comments from other authors

## Competing interests

The authors declare that the research was conducted in the absence of any commercial or financial relationships that could be construed as a potential conflict of interest.

## Notes

https://doi.org/10.6084/m9.figshare.21642209.v1

### Competing Interest Statement

The authors have declared no competing interest.

## References

1. Jones KE, Patel NG, Levy MA, Storeygard A, Balk D, Gittleman JL, et al. Global trends in emerging infectious diseases. Nature. 2008 Feb;451(4):990–3.

2. Hoelzer K, Shackelton LA, Holmes EC, Parrish CR. Within-Host Genetic Diversity of Endemic and Emerging Parvoviruses of Dogs and Cats. J Virol. 2008;82(22):11096–105.

3. Sharp PM, Hahn BH. The evolution of HIV-1 and the origin of AIDS. Philos Trans R Soc B Biol Sci. 2010;365(1552):2487–94.

4. Woolhouse MEJ, Adair K, Brierley L. RNA viruses : a case study of the biology of emerging infectious diseases. 2018;1(1):1–16.

5. Woolhouse M, Scott F, Hudson Z, Howey R, Chase-Topping M. Human viruses: discovery and emergence. Philos Trans R Soc London Ser B, Biol Sci. 2012 Oct;367(4):2864–71.

6. World Health Organization. HIV/AIDS. Global Health Observatory Data. 2018.

7. Johnson NPAS, Mueller J. Updating the accounts: global mortality of the 1918-1920 “Spanish” influenza pandemic. Bull Hist Med. 2002;76(1):105–15.

8. Murray CJ, Lopez AD, Chin B, Feehan D, Hill KH. Estimation of potential global pandemic influenza mortality on the basis of vital registry data from the 1918-20 pandemic: a quantitative analysis. Lancet. 2006;368(9554):2211–8.

9. Smith GJD, Vijaykrishna D, Bahl J, Lycett SJ, Worobey M, Pybus OG, et al. Origins and evolutionary genomics of the 2009 swine-origin H1N1 influenza a epidemic. Nature [Internet]. 2009;459(7250):1122–5. Available from: http://dx.doi.org/10.1038/nature08182

10. Worobey M, Han GZ, Rambaut A. A synchronized global sweep of the internal genes of modern avian influenza virus. Nature. 2014;508(7495):254–7.

11. Lytras S, Xia W, Hughes J, Jiang X, Robertson DL. The animal origin of SARS-CoV-2. Science (80-). 2021;373(6558):968–70.

12. Holmes EC, Goldstein SA, Rasmussen AL, Robertson DL, Crits-Christoph A, Wertheim JO, et al. The origins of SARS-CoV-2: A critical review. Cell [Internet]. 2021;184(19):4848–56. Available from: https://doi.org/10.1016/j.cell.2021.08.017

13. Andersen KG, Rambaut A, Lipkin WI, Holmes EC, Garry RF. The proximal origin of SARS-CoV-2. Vol. 26, Nature medicine. 2020. p. 450–2.

14. Guth S, Visher E, Boots M, Brook CE. Host phylogenetic distance drives trends in virus virulence and transmissibility across the animal-human interface. Philos Trans R Soc B Biol Sci. 2019;374(1782).

15. Farrell MJ, Davies TJ. Disease mortality in domesticated animals is predicted by host evolutionary relationships. Proc Natl Acad Sci U S A. 2019;116(16):7911–5.

16. Mollentze N, Streicker DG, Murcia PR, Hampson K, Biek R. Virulence mismatches in index hosts shape the outcomes of cross-species transmission. Proc Natl Acad Sci U S A. 2020;117(46):28859–66.

17. Longdon B, Hadfield JD, Day JP, Smith SCL, McGonigle JE, Cogni R, et al. The Causes and Consequences of Changes in Virulence following Pathogen Host Shifts. PLoS Pathog. 2015;11(3):1–18.

18. Perlman SJ, Jaenike J. Infection Success in Novel Hosts: An Experimental and Phylogenetic Study of Drosophila-parasitic Nematodes. Evolution (N Y). 2003;57(3):544–57.

19. Streicker DG, Turmelle AS, Vonhof MJ, Kuzmin I V, McCracken GF, Rupprecht CE. Host phylogeny constrains cross-species emergence and establishment of rabies virus in bats. Science. 2010 Aug;329(4):676–9.

20. Imrie RM, Roberts KE, Longdon B. Between virus correlations in the outcome of infection across host species: Evidence of virus by host species interactions. Evol Lett. 2021;5(5):472–83.

21. Longdon B, Hadfield JD, Webster CL, Obbard DJ, Jiggins FM. Host phylogeny determines viral persistence and replication in novel hosts. PLoS Pathog. 2011;7(9).

22. Tinsley MC, Majerus MEN. Small steps or giant leaps for male-killers? Phylogenetic constraints to male-killer host shifts. BMC Evol Biol. 2007;7(1):1–10.

23. Russell JA, Goldman-Huertas B, Moreau CS, Baldo L, Stahlhut JK, Werren JH, et al. Specialization and geographic isolation among Wolbachia symbionts from ants and lycaenid butterflies. Evolution [Internet]. 2008/11/19. 2009 Mar;63(4):624–40. Available from: https://www.ncbi.nlm.nih.gov/pubmed/19054050

24. Engelstädter J, Hurst GDD. The dynamics of parasite incidence across host species. Evol Ecol. 2006;20(6):603–16.

25. De Vienne DM, Hood ME, Giraud T. Phylogenetic determinants of potential host shifts in fungal pathogens. J Evol Biol. 2009;22(12):2532–41.

26. Gilbert GS, Webb CO. Phylogenetic signal in plant pathogen-host range. Proc Natl Acad Sci U S A. 2007;104(12):4979–83.

27. Faria NR, Suchard MA, Rambaut A, Streicker DG, Lemey P. Simultaneously reconstructing viral crossspecies transmission history and identifying the underlying constraints. Philos Trans R Soc B Biol Sci. 2013;368(1614).

28. Weinert LA, Welch JJ, Suchard MA, Lemey P, Rambaut A, Fitzgerald JR. Molecular dating of human-to-bovid host jumps by Staphylococcus aureus reveals an association with the spread of domestication. Biol Lett. 2012;8(5):829–32.

29. Webby RJ, Webster RG. Emergence of influenza A viruses. Philos Trans R Soc B Biol Sci. 2001;356(1416):1817–28.

30. Wille M, Lisovski S, Roshier D, Ferenczi M, Hoye BJ, Leen T, et al. Strong phylogenetic and ecological effects on host competency for avian influenza in Australian wild birds. bioRxiv [Internet]. 2022 Jan 1;2022.02.14.480463. Available from: http://biorxiv.org/content/early/2022/02/15/2022.02.14.480463.abstract

31. Barrow LN, McNew SM, Mitchell N, Galen SC, Lutz HL, Skeen H, et al. Deeply conserved susceptibility in a multi-host, multi-parasite system. Ecol Lett [Internet]. 2019 Jun 1;22(6):987–98. Available from: https://doi.org/10.1111/ele.13263

32. Martel A, Blooi M, Adriaensen C, Van Rooij P, Beukema W, Fisher MC, et al. Wildlife disease. Recent introduction of a chytrid fungus endangers Western Palearctic salamanders. Science. 2014 Oct;346(4):630–1.

33. Bernheim A, Sorek R. The pan-immune system of bacteria: antiviral defence as a community resource. Nat Rev Microbiol [Internet]. 2020;18(2):113–9. Available from: http://dx.doi.org/10.1038/s41579-019-0278-2

34. Dimitriu T, Marchant L, Buckling A, Raymond B. Bacteria from natural populations transfer plasmids mostly towards their kin. Proceedings Biol Sci. 2019 Jun;286(4):20191110.

35. Sakoparnig T, Field C, van Nimwegen E. Whole genome phylogenies reflect the distributions of recombination rates for many bacterial species. Nourmohammad A, Walczak AM, editors. Elife [Internet]. 2021;10:e65366. Available from: https://doi.org/10.7554/eLife.65366

36. World Health Organization. New report calls for urgent action to avert antimicrobial resistance crisis [Internet]. 2019. Available from: https://www.who.int/news-room/detail/29-04-2019-new-report-calls-for-urgent-action-to-avert-antimicrobial-resistance-crisis

37. AMR Industry Alliance. 2020 progress report [Internet]. 2020. Available from: https://www.amrindustryalliance.org/wp-content/uploads/2020/01/AMR-2020-Progress-Report.pdf

38. Perros M. A sustainable model for antibiotics. Science (80-) [Internet]. 2015;347(6226):1062–4. Available from: https://www.science.org/doi/10.1126/science.aaa3048

39. Bhargava K, Nath G, Bhargava A, Aseri GK, Jain N. Phage therapeutics: from promises to practices and prospectives. Appl Microbiol Biotechnol [Internet]. 2021;9047–67. Available from: https://doi.org/10.1007/s00253-021-11695-z

40. Fish R, Kutter E, Wheat G, Blasdel B, Kutateladze M, Kuhl S. Bacteriophage treatment of intransigent Diabetic toe ulcers: A case series. J Wound Care. 2016;25:S27–33.

41. Lin DM, Koskella B, Lin HC. Phage therapy: An alternative to antibiotics in the age of multi-drug resistance. World J Gastrointest Pharmacol Ther. 2017;8(3):162.

42. Van Nieuwenhuyse B, Van der Linden D, Chatzis O, Lood C, Wagemans J, Lavigne R, et al. Bacteriophage-antibiotic combination therapy against extensively drug-resistant Pseudomonas aeruginosa infection to allow liver transplantation in a toddler. Nat Commun [Internet]. 2022;13(1):5725. Available from: https://doi.org/10.1038/s41467-022-33294-w

43. Ross A, Ward S, Hyman P. More Is Better: Selecting for Broad Host Range Bacteriophages. Front Microbiol [Internet]. 2016;7:1352. Available from: https://www.frontiersin.org/article/10.3389/fmicb.2016.01352

44. Weinbauer MG. Ecology of prokaryotic viruses. FEMS Microbiol Rev. 2004;28(2):127–81.

45. Dekel-Bird NP, Sabehi G, Mosevitzky B, Lindell D. Host-dependent differences in abundance, composition and host range of cyanophages from the Red Sea. Environ Microbiol. 2015 Apr;17(4):1286–99.

46. Moebus K. Marine bacteriophage reproduction under nutrient-limited growth of host bacteria. II. Investigations with phage-host system [H3:H3/1]. Mar Ecol Prog Ser [Internet]. 1996;144:13–22. Available from: https://www.int-res.com/abstracts/meps/v144/p13-22/

47. Kauffman KM, Hussain FA, Yang J, Arevalo P, Brown JM, Chang WK, et al. A major lineage of non-tailed dsDNA viruses as unrecognized killers of marine bacteria. Nature. 2018;554(7690):118–22.

48. Munson-Mcgee JH, Peng S, Dewerff S, Stepanauskas R, Whitaker RJ, Weitz JS, et al. A virus or more in (nearly) every cell: Ubiquitous networks of virus-host interactions in extreme environments. ISME J [Internet]. 2018;12(7):1706–14. Available from: http://dx.doi.org/10.1038/s41396-018-0071-7

49. Duyvejonck H, Merabishvili M, Pirnay JP, De Vos D, Verbeken G, Van Belleghem J, et al. Development of a qPCR platform for quantification of the five bacteriophages within bacteriophage cocktail 2 (BFC2). Sci Rep [Internet]. 2019;9(1):1–10. Available from: http://dx.doi.org/10.1038/s41598-019-50461-0

50. Anderson B, Rashid MH, Carter C, Pasternack G, Rajanna C, Revazishvili T, et al. Enumeration of bacteriophage particles. Bacteriophage [Internet]. 2011;1(2):86–93. Available from: https://doi.org/10.4161/bact.1.2.15456

51. Khan Mirzaei M, Nilsson AS. Isolation of phages for phage therapy: a comparison of spot tests and efficiency of plating analyses for determination of host range and efficacy. PLoS One. 2015;10(3):e0118557.

52. Kwan T, Liu J, DuBow M, Gros P, Pelletier J. The complete genomes and proteomes of 27 Staphylococcus aureus bacteriophages. Proc Natl Acad Sci U S A [Internet]. 2005/03/23. 2005 Apr 5;102(14):5174–9. Available from: https://pubmed.ncbi.nlm.nih.gov/15788529

53. Vandersteegen K, Mattheus W, Ceyssens PJ, Bilocq F, de Vos D, Pirnay JP, et al. Microbiological and molecular assessment of bacteriophage ISP for the control of Staphylococcus aureus. PLoS One. 2011;6(9).

54. Synnott AJ, Kuang Y, Kurimoto M, Yamamichi K, Iwano H, Tanji Y. Isolation from sewage influent and characterization of novel Staphylococcus aureus bacteriophages with wide host ranges and potent lytic capabilities. Appl Environ Microbiol. 2009 Jul;75(4):4483–90.

55. Hsieh S-E, Lo H-H, Chen S-T, Lee M-C, Tseng Y-H. Wide host range and strong lytic activity of Staphylococcus aureus lytic phage Stau2. Appl Environ Microbiol. 2011 Feb;77(4):756–61.

56. Alves DR, Gaudion A, Bean JE, Perez Esteban P, Arnot TC, Harper DR, et al. Combined use of bacteriophage K and a novel bacteriophage to reduce Staphylococcus aureus biofilm formation. Appl Environ Microbiol. 2014 Nov;80(4):6694–703.

57. O’Flaherty S, Ross RP, Meaney W, Fitzgerald GF, Elbreki MF, Coffey A. Potential of the polyvalent anti-Staphylococcus bacteriophage K for control of antibiotic-resistant staphylococci from hospitals. Appl Environ Microbiol. 2005 Apr;71(4):1836–42.

58. Gutiérrez D, Briers Y, Rodríguez-Rubio L, Martínez B, Rodríguez A, Lavigne R, et al. Role of the Pre-neck Appendage Protein (Dpo7) from Phage vB_SepiS-phiIPLA7 as an Anti-biofilm Agent in Staphylococcal Species. Front Microbiol. 2015;6:1315.

59. Verstappen KM, Tulinski P, Duim B, Fluit AC, Carney J, van Nes A, et al. The Effectiveness of Bacteriophages against Methicillin-Resistant Staphylococcus aureus ST398 Nasal Colonization in Pigs. PLoS One. 2016;11(8):e0160242.

60. Wills QF, Kerrigan C, Soothill JS. Experimental bacteriophage protection against Staphylococcus aureus abscesses in a rabbit model. Antimicrob Agents Chemother. 2005 Mar;49(4):1220–1.

61. Matsuzaki S, Yasuda M, Nishikawa H, Kuroda M, Ujihara T, Shuin T, et al. Experimental protection of mice against lethal Staphylococcus aureus infection by novel bacteriophage phi MR11. J Infect Dis. 2003 Feb;187(4):613–24.

62. Chatterjee AN, Mirelman D, Singer HJ, Park JT. Properties of a novel pleiotropic bacteriophage-resistant mutant of Staphylococcus aureus H. J Bacteriol. 1969 Nov;100(4):846–53.

63. Chatterjee AN. Use of bacteriophage-resistant mutants to study the nature of the bacteriophage receptor site of Staphylococcus aureus. J Bacteriol. 1969 May;98(4):519–27.

64. Shaw DR, Chatterjee AN. O-Acetyl groups as a component of the bacteriophage receptor on Staphylococcus aureus cell walls. J Bacteriol. 1971 Oct;108(4):584–5.

65. Shaw DR, Mirelman D, Chatterjee AN, Park JT. Ribitol teichoic acid synthesis in bacteriophage-resistant mutants of Staphylococcus aureus H. J Biol Chem. 1970 Oct;245(4):5101–6.

66. Brown S, Santa Maria JPJ, Walker S. Wall teichoic acids of gram-positive bacteria. Annu Rev Microbiol. 2013;67:313–36.

67. Xia G, Corrigan RM, Winstel V, Goerke C, Gründling A, Peschel A. Wall teichoic Acid-dependent adsorption of staphylococcal siphovirus and myovirus. J Bacteriol. 2011 Aug;193(4):4006–9.

68. Wilkinson BJ, Holmes KM. Staphylococcus aureus cell surface: capsule as a barrier to bacteriophage adsorption. Infect Immun. 1979 Feb;23(4):549–52.

69. Ohshima Y, Schumacher-Perdreau F, Peters G, Pulverer G. The role of capsule as a barrier to bacteriophage adsorption in an encapsulated Staphylococcus simulans strain. Med Microbiol Immunol. 1988;177(4):229–33.

70. Nordström K, Forsgren A. Effect of protein A on adsorption of bacteriophages to Staphylococcus aureus. J Virol. 1974;14(2):198–202.

71. Sadykov MR. Restriction–modification systems as a barrier for genetic manipulation of Staphylococcus aureus. In: The genetic manipulation of staphylococci. Springer; 2014. p. 9–23.

72. Monk IR, Foster TJ. Genetic manipulation of Staphylococci-breaking through the barrier. Front Cell Infect Microbiol. 2012;2:49.

73. Costa SK, Donegan NP, Corvaglia A-R, François P, Cheung AL. Bypassing the Restriction System To Improve Transformation of Staphylococcus epidermidis. J Bacteriol. 2017 Aug;199(16).

74. Yang S, Liu J, Shao F, Wang P, Duan G, Yang H. Analysis of the features of 45 identified CRISPR loci in 32 Staphylococcus aureus. Biochem Biophys Res Commun. 2015 Aug;464(4):894–900.

75. Li Q, Xie X, Yin K, Tang Y, Zhou X, Chen Y, et al. Characterization of CRISPR-Cas system in clinical Staphylococcus epidermidis strains revealed its potential association with bacterial infection sites. Microbiol Res. 2016 Dec;193:103–10.

76. Barrangou R, Fremaux C, Deveau H, Richards M, Boyaval P, Moineau S, et al. CRISPR provides acquired resistance against viruses in prokaryotes. Science. 2007 Mar;315(4):1709–12.

77. Barrangou R, Marraffini LA. CRISPR-Cas systems: Prokaryotes upgrade to adaptive immunity. Mol Cell. 2014 Apr;54(4):234–44.

78. Kumar S, Suleski M, Craig JM, Kasprowicz AE, Sanderford M, Li M, et al. TimeTree 5: An Expanded Resource for Species Divergence Times. Mol Biol Evol [Internet]. 2022 Aug 1;39(8):msac174. Available from: https://doi.org/10.1093/molbev/msac174

79. Ruijter JM, Thygesen HH, Schoneveld OJLM, Das AT, Berkhout B, Lamers WH. Factor correction as a tool to eliminate between-session variation in replicate experiments: Application to molecular biology and retrovirology. Retrovirology. 2006;3(May).

80. Ruijter JM, Ruiz Villalba A, Hellemans J, Untergasser A, van den Hoff MJB. Removal of between-run variation in a multi-plate qPCR experiment. Biomol Detect Quantif [Internet]. 2015;5:10–4. Available from: http://dx.doi.org/10.1016/j.bdq.2015.07.001

81. Drummond AJ, Suchard MA, Xie D, Rambaut A. Bayesian phylogenetics with BEAUti and the BEAST 1.7. Mol Biol Evol. 2012 Aug;29(4):1969–73.

82. Seemann T. Prokka: Rapid prokaryotic genome annotation. Bioinformatics. 2014 Jul;30(4):2068–9.

83. Tonkin-Hill G, MacAlasdair N, Ruis C, Weimann A, Horesh G, Lees JA, et al. Producing polished prokaryotic pangenomes with the Panaroo pipeline. Genome Biol. 2020 Jul;21(4):1–21.

84. Shapiro B, Rambaut A, Drummond AJ. Choosing appropriate substitution models for the phylogenetic analysis of protein-coding sequences. Vol. 23, Molecular biology and evolution. United States; 2006. p. 7–9.

85. Rambaut A, Suchard M, Xie D, Drummond A. Tracer v1.6 [Internet]. 2014. Available from: http://beast.bio.ed.ac.uk/Tracer

86. Paradis E, Schliep K. ape 5.0: an environment for modern phylogenetics and evolutionary analyses in R. Bioinformatics [Internet]. 2019 Feb 1;35(3):526–8. Available from: https://doi.org/10.1093/bioinformatics/bty633

87. Hadfield JD. MCMC Methods for Multi-Response Generalized Linear Mixed Models: The MCMCglmm R Package. J Stat Softw [Internet]. 2010 Feb 2;33(2 SE-Articles):1–22. Available from: https://www.jstatsoft.org/index.php/jss/article/view/v033i02

88. Freckleton RP, Harvey PH, Pagel M. Phylogenetic Analysis and Comparative Data: A Test and Review of Evidence. Am Nat [Internet]. 2002 Dec 1;160(6):712–26. Available from: https://doi.org/10.1086/343873

89. Housworth EA, Martins EP, Lynch M. The Phylogenetic Mixed Model. Am Nat. 2004;163(1):84–96.

90. Falconer DS. Introduction to quantitative genetics. Pearson Education India; 1996.

91. Poulin R, Krasnov BR, Mouillot D, Thieltges DW. The comparative ecology and biogeography of parasites. Philos Trans R Soc B Biol Sci [Internet]. 2011 Aug 27;366(1576):2379–90. Available from: https://doi.org/10.1098/rstb.2011.0048

92. de Jonge PA, Nobrega FL, Brouns SJJ, Dutilh BE. Molecular and Evolutionary Determinants of Bacteriophage Host Range. Trends Microbiol. 2019 Jan;27(4):51–63.

93. Chow LT, Bukhari AI. The invertible DNA segments of coliphages Mu and P1 are identical. Virology [Internet]. 1976 Oct;74(4):242—248. Available from: https://doi.org/10.1016/0042-6822(76)90148-3

94. Liu M, Deora R, Doulatov SR, Gingery M, Eiserling FA, Preston A, et al. Reverse transcriptase-mediated tropism switching in Bordetella bacteriophage. Science. 2002 Mar;295(4):2091–4.

95. Meyer JR, Dobias DT, Weitz JS, Barrick JE, Quick RT, Lenski RE. Repeatability and Contingency in the Evolution of a Key Innovation in Phage Lambda. Science (80-) [Internet]. 2012;335(6067):428–32. Available from: https://science.sciencemag.org/content/335/6067/428

96. Takeuchi I, Osada K, Azam AH, Asakawa H, Miyanaga K, Tanji Y. The Presence of Two Receptor-Binding Proteins Contributes to the Wide Host Range of Staphylococcal Twort-Like Phages. Appl Environ Microbiol. 2016 Oct;82(4):5763–74.

97. Moller AG, Lindsay JA, Read TD. Determinants of Phage Host Range in Staphylococcus Species. Appl Environ Microbiol. 2019 Jun;85(11).

98. Koonin E V, Makarova KS, Wolf YI. Evolutionary Genomics of Defense Systems in Archaea and Bacteria. Annu Rev Microbiol. 2017 Sep;71:233–61.

99. Oliveira PH, Touchon M, Rocha EPC. The interplay of restriction-modification systems with mobile genetic elements and their prokaryotic hosts. Nucleic Acids Res. 2014;42(16):10618–31.

100. Makarova KS, Wolf YI, Alkhnbashi OS, Costa F, Shah SA, Saunders SJ, et al. An updated evolutionary classification of CRISPR–Cas systems. Nat Rev Microbiol [Internet]. 2015;13(11):722–36. Available from: https://doi.org/10.1038/nrmicro3569

101. Godde JS, Bickerton A. The repetitive DNA elements called CRISPRs and their associated genes: evidence of horizontal transfer among prokaryotes. J Mol Evol. 2006 Jun;62(4):718–29.

102. Makarova KS, Wolf YI, Koonin E V. Comparative genomics of defense systems in archaea and bacteria. Nucleic Acids Res. 2013 Apr;41(4):4360–77.

103. Abedon ST. Lysis from without. Bacteriophage. 2011 Jan;1(4):46–9.

104. Secor PR, James GA, Fleckman P, Olerud JE, McInnerney K, Stewart PS. Staphylococcus aureus Biofilm and Planktonic cultures differentially impact gene expression, mapk phosphorylation, and cytokine production in human keratinocytes. BMC Microbiol [Internet]. 2011;11(1):143. Available from: http://www.biomedcentral.com/1471-2180/11/143

105. Otto M. Staphylococcal Biofilms. Microbiol Spectr. 2018 Aug;6(4).

106. Caiazza NC, O’Toole GA. Alpha-toxin is required for biofilm formation by Staphylococcus aureus. J Bacteriol. 2003 May;185(4):3214–7.

107. Resch A, Rosenstein R, Nerz C, Götz F. Differential gene expression profiling of Staphylococcus aureus cultivated under biofilm and planktonic conditions. Appl Environ Microbiol. 2005 May;71(4):2663–76.

