## Supplementary Text for "The host phylogeny determines viral infectivity and replication across *Staphylococcus* host species"

Complete supplementary materials

**Running title: Phylogenetic determinants of phage host range**

Sarah K Walsh[1,2*], Ryan M Imrie[1], Marta Matuszewska[3], Gavin K Paterson[4], Lucy A Weinert[5], Jarrod D Hadfield[6], Angus Buckling[1,2+], and Ben Longdon[1+]

[1]Centre for Ecology and Conservation; Faculty of Environment, Science, and Economy; Biosciences; University of Exeter; Penryn Campus; Penryn; Cornwall; TR10 9FE; UK.

[2]Environment and Sustainability Institute; University of Exeter; Penryn Campus; Penryn; TR10 9FE; Cornwall; UK.

[3]Department of Medicine; University of Cambridge; Cambridge; CB2 0SP; UK.

[4]Royal (Dick) School of Veterinary Studies and the Roslin Institute; University of Edinburgh; Easter Bush; Midlothian; EH25 9RG; UK.

[5]Department of Veterinary Medicine; University of Cambridge; Cambridge; CB3 0ES; UK.

[6]Institute of Evolutionary Biology; School of Biological Sciences; The University of Edinburgh; Edinburgh; EH9 3JT; UK.

[+] Equal contribution

### Contents

1. Supplementary Methods
    - 1.1. Sample preparation and DNA extraction
    - 1.2. Illumina sequencing
    - 1.3. Whole genome sequence quality control and *de novo* assembly
    - 1.4. Construction of a phylogenetic tree for the 48 *S. aureus* strains
    - 1.5. Construction of a core genome alignment for the *Staphylococcus* host panel
    - 1.6. Construction of an ultrametric phylogenetic tree for the *Staphylococcus* host panel
2. Supplementary Tables
    - 2.1. Supplementary Table 1: Staphylococcus metadata and sequencing data.
    - 2.2. Supplementary Table 2: Estimates for the repeatability and heritability across 64 *Staphylococcus* isolates when no outliers were removed from the OD data.
    - 2.3. Supplementary Table 3: Estimates for the repeatability and heritability across 64 Staphylococcus isolates when major outliers were removed from the OD data.
    - 2.4. Supplementary Table 4: Estimates from the phylogenetic generalised linear mixed models for inter-strain correlations in susceptibility between methods where no outliers were removed from the OD data.
    - 2.5. Supplementary Table 5: Estimates from the phylogenetic generalised linear mixed models for inter-strain correlations in susceptibility between methods where major outliers were removed from the OD data.
    - 2.6. Supplementary Table 6: Estimates from the S. aureus only model for the repeatability and heritability.
    - 2.7. Supplementary Table 7: Estimates from the species only model for repeatability and heritability.
    - 2.8. Supplementary Table 8: Inter-strain correlations between methods for assessing host range in a within-aureus model.
    - 2.9. Supplementary Table 9: Inter-species correlations between methods for assessing host range in aa among-species model.
    - 2.10. Table S10: *Staphylococcus* sequences used in the FastQ Screen to determine if all sequences used in this study belong to *Staphylococcus*/*S. aureus*.
3. Supplementary Figures
    - 3.1. Supplementary Figure 1: A comparison of all 123 trees on 64 tips using the Kendall Colijn metric vector (mid-point rooted gene trees).
4. Supplementary References

1. Supplementary Methods

### 1.1. Sample preparation and DNA extraction

To infer the relationship between host species, 56 of the *Staphylococcus* isolates were whole genome sequenced. Pure cultures of each of the strains were grown on LB agar plates at 37$^{o}$C overnight. Bacteria were then harvested from the plates and resuspended in a 2mL tube containing cryoperservative (Microbank$^{TM}$, Pro-Lab Diagnostics UK, United Kingdom) according to MicrobesNG instructions. Samples were then shipped to MicrobesNG where they were processed as follows: 5-40uL of the bacterial suspension were lysed with 120uL of TE buffer containing lysozyme (final concentration 0.1mg/mL) and RNase A (ITW Reagents, Barcelona, Spain) (final concentration 0.1mg/mL) and incubated for 25 minutes at 37$^{o}$C. Proteinase K (VWR Chemicals, Ohio, USA) (final concentration 0.1mg/mL) and SDS (Sigma-Aldrich, Missouri, USA) (final concentration 0.5% v/v) are added and incubated for 5 minutes at 65$^{o}$C. Genomic DNA was then purified using an equal volume of SPRI beads and resuspended in EB buffer (Qiagen, Germany). DNA was quantified using a Quant-iT dsDNA HS kit (ThermoFisher Scientifitc) assay in an Eppendorf AF2200 plate reader (Eppendorf UK Ltd, United Kingdom).

### 1.2. Illumina sequencing

Library preparation and sequencing were performed by MicrobesNG (Birmingham, UK). Briefly, genomic DNA libraries were prepared using the Nextera XT Library Prep Kit (Illumina, San Diego, USA) with the following modifications: twice the amount of input DNA is used, and PCR elongation is increased to 45-seconds. Pooled libraries were quantified using the Kapa Biosystems Library Quantification Kit for Illumina and sequenced using Illumina sequencers (HiSeq/NovaSeq) with a 25-bp paired end protocol.

### 1.3. Whole genome sequence quality control and *de novo* assembly

To supplement the 56 samples that were sequenced by MicrobesNG, the whole genome sequences of the remaining 8 samples were downloaded from NCBI (Supplementary Table 1). Adapters and low-quality reads were removed with *Cutadapt* v1.16 (1) and *Sickle* v1.33 (2) and reads screened for contamination using FastQ Screen v0.12.0 (3). *De novo* assembly was performed using the SPAdes genome assembler (v3.9.1) (4). Optimal k-mers were identified based on the average read lengths for each genome.

All assemblies were evaluated using QUAST v.5.0.2 (5) and reads mapped back to *de novo* assemblies to identify potential contamination using *Bowtie2* v2.3.4.1 (6). Low-quality genome assemblies were excluded from further analysis, defined in the following way: all assemblies with an N50 of less than 10kb; *S. aureus* total sequence length plus or minus one standard deviation outside of the median sequence length based on previously collected data of 2011 *S. aureus* isolates (the median and standard deviation of genome size for other *Staphylococcal* species was evaluated based on published genomes); all assemblies with greater than 1500 SNPs (this threshold was defined by identifying a distribution of SNPs in all assemblies); and finally, if the total length of contigs smaller than 1kb exceeded 15% of the total genome assembly length (often indicative of a sample with >1 bacterial isolate), but only if one or more other QC measure was not met.

A FastQ Screen v0.12.0 was used to check post-sickle reads for contamination and to determine the identification of a *Staphylococcus* species. This was further confirmed via Kraken (3,7) by creating a custom database of *Staphylococcus* reference genomes (Table S10) along with a built-in database with PhiX of vectors and contaminants commonly seen in sequencing experiments (3). The custom database was created by downloading the *Staphylococcus* reference genome sequences in FASTA format from NCBI and then using Bowtie2 to build the relevant index files (6). Contaminants were detected by comparing the percentage of reads mapped uniquely against each of the reference genomes and additionally investigated using Kraken. Contamination was assumed if the reads mapped to a reference strain <70% and mapped to other species >30%. Ten isolates were identified in FastQ screen below >70% match and nine of these were <70% in Kraken. 271Y was mapped to *Staphylococcus kloosii* at 67.38% but this specific species only has 18 genomes available on NCBI, which most likely does not represent the species diversity and as other QC measures were within acceptable range was not excluded.

#### 1.4. Construction of a phylogenetic tree for the 48 *S. aureus* strains

For the phylogenetic tree containing only the *S. aureus* isolates, reference-mapped assemblies were generated using Bowtie2 v1.2.2. In order to identify a reference strain outside of the collection diversity, sequence types (STs) for each *S. aureus* genome were identified using MLST-check (8). The reference genome S0385 was selected as none of the *S. aureus* were identified as ST398 (GenBank accession no. AM990992) (6,9). In order to map completed reference genomes, artificial FASTQ files were generated using ArtificialFastqGenerator (10).

Recombination was identified in the reference-mapped alignment using both Gubbins v2.3.1 (11) and ClonalFrame (12). All identified recombinant sites were masked, including a previously identified ~123 kb genome segment horizontally acquired from an ST9 donor (13).

Phylogenetic reconstruction was carried out for the reference-mapped alignment with RAxML (v8.2.4) using the GTR+Γ model and 1,000 bootstraps (14) with a *Staphylococcus argenteus* outgroup (MSHR1132) to establish the root. The phylogenetic reconstruction was then carried out excluding the outgroup. To only include phylogenetically informative sites, sites containing more than a single gap were excluded due to marked recombination or missing data. Trees were visualized and annotated with Figtree (http://tree.bio.ed.ac.uk/software/figtree/) and the Interactive Tree Of Life (15).

#### 1.5. Construction of a core genome alignment for the *Staphylococcus* host panel

D*e novo* assemblies were annotated using Prokka (v2.8.2) (16) and orthologous genes identified with Panaroo (17) using a range of different sequence identity thresholds (0.95-0.70). A sequence identity threshold of 0.7 was selected to maximise the number of orthologous groups with only one representative in each of the 64 isolates, while reducing the homology groups containing duplicates. This threshold identified 129 genes, of which six were excluded due non-orthologous groups or having at least a single gap within all the sites in the gene alignment. Each of the remaining 123 identified orthologous groups was investigated by creating a single gene neighbour joining tree using the ape package in R and a K80 substitution model (18). Contradictory gene topologies were identified using the R package “treescape” (19) as previously described by (20) using a multi-dimensional scaling visualization of tree distances (Figure S1). From this, 21 of the genes with the most district tree topologies were excluded. The final core genome alignment was created based on 102 genes (125,893 bp).

#### 1.6. Construction of an ultrametric phylogenetic tree for the *Staphylococcus* host panel

An ultrametric phylogenetic tree was constructed using BEAST v1.10 with an HKY+Γ model, an uncorrelated relaxed molecular clock, and constant population size coalescent prior, from the reference-mapped alignment for *S. aureus* and core genome for all *Staphylococc*i spp. (Drummond et al., 2012). For the *S. aureus* reference-mapped phylogenetic reconstruction I fitted separate substitution models and molecular clocks to 1st/2nd, and 3rd codon positions, non-coding and RNA positions to reflect differences in selective constraint (21). For the core genome Staphylococci phylogenetic reconstruction, we fitted separate substitution models and molecular clocks to 1st/2nd, and 3rd codon positions, to reflect differences in selective constraint (21). For the two models, we ran two independent MCMC until convergence was reached and the burn-in represented <10% of the chain. Convergence of all parameters was checked in the program Tracer v1.4 (22).

2. Supplementary Tables

**Supplementary Table 1: Staphylococcus metadata and sequencing data**.

| Isolate ID | Species | Host | Location | Year | NCBI BioSample ID | Obtained from |
|---|---|---|---|---|---|---|
| 13S44S9 | *S. aureus* | *Homo sapiens* | BEL | 2012 | SAMN31484028 | Jean-Paul Pirnay |
| 271Y | *S. kloosii* | *Eptesicus serotinus* | GBR | 2015 | SAMN31484029 | Edward Feil |
| 2111F7LW | *S. sciuri* | *Pteropus livingstonii* | GBR | 2017 | SAMN31484030 | Edward Feil |
| 27420LC | *S. simiae* | *Pteropus livingstonii* | GBR | 2016 | SAMN31484031 | Edward Feil |
| 2745SW | *S. nepalensis* | *Pteropus livingstonii* | GBR | 2016 | SAMN31484032 | Edward Feil |
| 82B | *S. caeli* | Environmental | ITA | 2013 | SAMEA2297795 | Gavin Paterson |
| 8325-4 | *S. aureus* | --- | --- | --- | SAMN31484033 | Edward Feil |
| AR03918O1 | *S. aureus* | *Sciurus carolinensis* | GBR | 2018 | SAMN31484034 | Edward Feil |
| AR05S1 | *S. aureus* | *Sciurus carolinensis* | GBR | 2015 | SAMN31484035 | Edward Feil |
| AR05618O1 | *S. aureus* | *Sciurus carolinensis* | GBR | 2018 | SAMN31484036 | Edward Feil |
| ASARM61 | *S. aureus* | *Homo sapiens* | GBR | 2006 | SAMN31484037 | Edward Feil |
| ASARM70 | *S. aureus* | *Homo sapiens* | GBR | 2006 | SAMN31484038 | Edward Feil |
| ASARM71 | *S. aureus* | *Homo sapiens* | GBR | 2006 | SAMN31484039 | Edward Feil |
| ASARM72 | *S. aureus* | *Homo sapiens* | GBR | 2006 | SAMN31484040 | Edward Feil |
| ASARM73 | *S. aureus* | *Homo sapiens* | GBR | 2006 | SAMN31484041 | Edward Feil |
| ASARM74 | *S. aureus* | *Homo sapiens* | GBR | 2006 | SAMN31484042 | Edward Feil |
| NCTC7692 | *S. saprophyticus spp saprophyticus* | Environmental | --- | 1948 | SAMEA3517999 | Gavin Paterson |
| NCTC11320 | *S. hominis spp hominis* | *Homo sapiens* | USA | 1975 | SAMEA3539708 | Gavin Paterson |
| NCTC11043 | *S. xylosus* | *Homo sapiens* | USA | 1975 | SAMEA3539705 | Gavin Paterson |
| B128S3 | *S. aureus* | *Sciurus carolinensis* | GBR | 2015 | SAMN31484043 | Edward Feil |
| B142S1 | *S. aureus* | *Sciurus carolinensis* | GBR | 2015 | SAMN31484044 | Edward Feil |

| | | | | | | |
|---|---|---|---|---|---|---|
| DAR04181C1 | *S. aureus* | *Cervus elaphus* | GBR | 2018 | SAMN31484045 | Edward Feil |
| DAR06181LC1 | *S. aureus* | *Cervus elaphus* | GBR | 2018 | SAMN31484046 | Edward Feil |
| DAR091813 | *S. aureus* | *Cervus elaphus* | GBR | 2018 | SAMN31484047 | Edward Feil |
| DEU1 | *S. aureus* | *Homo sapiens* | TUR | 2009 | SAMN31484048 | Edward Feil |
| DEU2 | *S. aureus* | *Homo sapiens* | TUR | 2009 | SAMN31484049 | Edward Feil |
| DSM10441 | *S. edaphicus* | Environmental | ATA | 2013 | SAMN31484050 | Gavin Paterson |
| DSM107950 | *S. pseudoxylosus* | *Bos taurus* | FRA | 2002 | SAMN31484051 | Gavin Paterson |
| DSM18669 | *S. saprophyticus spp Bovis* | *Bos taurus* | CZE | 1996 | SAMN31484052 | Gavin Paterson |
| DSM21284 | *S. pseudointermedius* | *Felis catus* | BEL | 2008 | SAMN31484053 | Gavin Paterson |
| NCTC12218 | *S. schleiferi spp coagulans* | *Homo sapiens* | --- | 1988 | SAMEA3221103 | Gavin Paterson |
| DSM6628 | *S. schleiferi spp schleiferi* | *Canis lupus* | --- | 1991 | SAMN31484054 | Gavin Paterson |
| EOE23 | *S. aureus* | *Homo sapiens* | GBR | 1998 | SAMN31484055 | Edward Feil |
| EOE03 | *S. aureus* | *Homo sapiens* | GBR | 1998 | SAMN31484056 | Edward Feil |
| EOE30 | *S. aureus* | *Homo sapiens* | GBR | 1998 | SAMN31484057 | Edward Feil |
| EOE35 | *S. aureus* | *Homo sapiens* | GBR | 2003 | SAMN31484058 | Edward Feil |
| EOE41 | *S. aureus* | *Homo sapiens* | GBR | 2005 | SAMN31484059 | Edward Feil |
| EOE42 | *S. aureus* | *Homo sapiens* | GBR | 2005 | SAMN31484060 | Edward Feil |
| HU25 | *S. aureus* | *Homo sapiens* | BRA | 1905 | SAMN31484061 | Edward Feil |
| JW32660O5 | *S. aureus* | *Sciurus carolinensis* | GBR | 2018 | SAMN31484062 | Edward Feil |
| JW30866OBHY3 | *S. aureus* | *Sciurus carolinensis* | GBR | 2018 | SAMN31484063 | Edward Feil |
| JW31330LBHY2 | *S. aureus* | *Sciurus carolinensis* | GBR | 2018 | SAMN31484064 | Edward Feil |
| JW31330OBHY1 | *S. aureus* | *Sciurus carolinensis* | GBR | 2018 | SAMN31484065 | Edward Feil |
| MU1 | *S. aureus* | *Homo sapiens* | TUR | 2010 | SAMN31484066 | Edward Feil |
| MU2 | *S. aureus* | *Homo sapiens* | TUR | 2010 | SAMN31484067 | Edward Feil |
| NCTC11042 | *S. haemolyticus* | *Homo sapiens* | CZE | 1976 | SAMEA3233544 | Gavin Paterson |

| | | | | | | |
|---|---|---|---|---|---|---|
| NCTC11046 | *S. simulans* | *Homo sapiens* | CZE | 1976 | SAMEA3504572 | Gavin Paterson |
| NCTC11047 | *S. epidermidis* | *Homo sapiens* | CZE | 1976 | SAMEA3233545 | Gavin Paterson |
| P32 | *S. aureus* | *Homo sapiens* | POL | 1996 | SAMN31484068 | Edward Feil |
| SaTPS3026 | *S. aureus* | *Homo sapiens* | AUS | 2013 | SAMN31484069 | Edward Feil |
| SaTPS3043 | *S. aureus* | *Homo sapiens* | AUS | 2013 | SAMN31484070 | Edward Feil |
| SaTPS3072 | *S. aureus* | *Homo sapiens* | AUS | 2013 | SAMN31484071 | Edward Feil |
| SaTPS3097 | *S. aureus* | *Homo sapiens* | AUS | 2013 | SAMN31484072 | Edward Feil |
| SaTPS3104 | *S. aureus* | *Homo sapiens* | AUS | 2013 | SAMN31484073 | Edward Feil |
| SaTPS3105 | *S. aureus* | *Homo sapiens* | AUS | 2013 | SAMN31484074 | Edward Feil |
| SAR1018S1 | *S. aureus* | *Ovis aries* | GBR | 2018 | SAMN31484075 | Edward Feil |
| SAR1218N1 | *S. aureus* | *Ovis aries* | GBR | 2018 | SAMN31484076 | Edward Feil |
| SAR1418N1 | *S. aureus* | *Ovis aries* | GBR | 2018 | SAMN31484077 | Edward Feil |
| USFL008 | *S. aureus* | *Homo sapiens* | USA | 2009 | SAMN31484078 | Edward Feil |
| USFL009 | *S. aureus* | *Homo sapiens* | USA | 2009 | SAMN31484079 | Edward Feil |
| USFL012 | *S. aureus* | *Homo sapiens* | USA | 2009 | SAMN31484080 | Edward Feil |
| USFL016 | *S. aureus* | *Homo sapiens* | USA | 2009 | SAMN31484081 | Edward Feil |
| USFL018 | *S. aureus* | *Homo sapiens* | USA | 2009 | SAMN31484082 | Edward Feil |
| USFL020 | *S. aureus* | *Homo sapiens* | USA | 2009 | SAMN31484083 | Edward Feil |

**Supplementary Table 2: Estimates for the repeatability and heritability across 64 *Staphylococcus* isolates when no outliers were removed from the OD data.** Estimates of repeatability are taken from model (2) and estimates of phylogenetic heritability (the variation explained by the host phylogeny) are taken from model (1). * indicates the phylogenetic heritability calculated as the proportion of variation that is attributed to phylogeny divided by the total variation (phylogenetic, non-phylogenetic between-strain, and within-strain variation) as opposed to phylogenetic and non-phylogenetic between-strain variation. PA = plaque assay, CI = credible interval.

| | **Repeatability** | | **Phylogenetic heritability** | | **Phylogenetic heritability of total variance*** | |
|---|---|---|---|---|---|---|
| **Method** | Mean | 95% CI | Mean | 95% CI | Mean | 95% CI |
| **Binary PA** | 1.00 | 1.00, 1.00 | 1.00 | 1.00, 1.00 | 1.00 | 1.00, 1.00 |
| **Continuous PA** | 0.00 | 0.00, 0.00 | 0.65 | 0.01, 1.00 | $9.93\times10^{-7}$ | $1.32\times10^{-13}$, $2.59\times10^{-6}$ |
| **OD** | 0.94 | 0.89, 0.98 | 0.99 | 0.96, 1.00 | 0.91 | 0.79, 0.98 |
| **qPCR** | 0.99 | 0.99, 0.99 | 1.00 | 1.00, 1.00 | 0.99 | 0.99, 1.00 |

**Supplementary Table 3: Estimates for the repeatability and heritability across 64 *Staphylococcus* isolates when major outliers were removed from the OD data.** Estimates of repeatability are taken from model (2) and estimates of phylogenetic heritability (the variation explained by the host phylogeny) are taken from model (1). * indicates the phylogenetic heritability calculated as the proportion of variation that is attributed to phylogeny divided by the total variation (phylogenetic, non-phylogenetic between-strain, and within-strain variation) as opposed to phylogenetic and non-phylogenetic between-strain variation. PA = plaque assay, CI = credible interval.

| | **Repeatability** | | **Phylogenetic heritability** | | **Phylogenetic heritability of total variance*** | |
|---|---|---|---|---|---|---|
| **Method** | Mean | 95% CI | Mean | 95% CI | Mean | 95% CI |
| **Binary PA** | 1.00 | 1.00, 1.00 | 1.00 | 1.00, 1.00 | 1.00 | 1.00, 1.00 |
| **Continuous PA** | 0.00 | 0.00, 0.00 | 0.68 | 0.01, 1.00 | $1.21\times10^{-6}$ | $1.56\times10^{-15}$, $5.20\times10^{-6}$ |
| **OD** | 0.98 | 0.96, 0.99 | 0.99 | 0.97, 1.00 | 0.95 | 0.91, 0.99 |
| **qPCR** | 0.99 | 0.99, 1.00 | 1.00 | 1.00, 1.00 | 0.99 | 0.98, 1.00 |

**Supplementary Table 4: Estimates from the phylogenetic generalised linear mixed models for inter-strain correlations in susceptibility between methods where no outliers were removed from the OD data.** Numbers show the mean estimates for the correlation strength (r, white cells) and slope (β, grey cells) between pairs of methods, with 95% credible intervals (CIs) indicated in brackets. The correlations and slopes were calculated with columns as x and rows as y. Estimates with CIs that do not span zero are highlighted in bold. PA = plaque assay, *value on a probit scale.

| | Binary PA | Continuous PA | OD | qPCR |
|---|---|---|---|---|
| **Binary PA** | - | - | **0.89 (0.75, 0.98)** | **0.97 (0.91, 1.00)** |
| **Continuous PA** | - | - | -0.00 (-0.97, 0.97) | -0.01 (-0.99, 1.00) |
| **OD** | **0.50* (0.50, 0.50)** | 0.02 (-0.33, 0.30) | - | **0.92 (0.83, 0.98)** |
| **qPCR** | **0.51* (0.51, 0.52)** | 0.23 (-1.77, 1.45) | **4.67 (3.20, 6.35)** | - |

**Supplementary Table 5: Estimates from the phylogenetic generalised linear mixed models for inter-strain correlations in susceptibility between methods where major outliers were removed from the OD data.** Numbers show the mean estimates for the correlation strength (r, white cells) and slope (β, grey cells) between pairs of methods, with 95% credible intervals (CIs) indicated in brackets. The correlations and slopes were calculated with columns as x and rows as y. Estimates with CIs that do not span zero are highlighted in bold. PA = plaque assay, *value on a probit scale.

| | Binary PA | Continuous PA | OD | qPCR |
|---|---|---|---|---|
| **Binary PA** | - | - | **0.95 (0.89, 0.99)** | **0.95 (0.95, 1.00)** |
| **Continuous PA** | - | - | -0.01 (-0.99, 0.99) | -0.01 (-1.00, 1.00) |
| **OD** | **0.50* (0.50, 0.50)** | 0.10 (-0.16, 0.23) | - | **0.97 (0.93, 1.00)** |
| **qPCR** | **0.51* (0.50, 0.52)** | 0.57 (-0.89, 1.20) | **5.26 (4.03, 6.55)** | - |

**Supplementary Table 6: Estimates from the *S. aureus* only model for the repeatability and heritability.** Estimates of repeatability are taken from model (2) and estimates of phylogenetic heritability (the variation explained by the host phylogeny) are taken from model (1). * indicates the phylogenetic heritability calculated as the proportion of variation that is attributed to phylogeny divided by the total variation (phylogenetic, non-phylogenetic between-strain, and within-strain variation) as opposed to phylogenetic and non-phylogenetic between-strain variation. PA = plaque assay, CI = credible interval.

| | **Repeatability** | | **Phylogenetic heritability** | | **Phylogenetic heritability of total variance*** | |
|---|---|---|---|---|---|---|
| **Method** | Mean | 95% CI | Mean | 95% CI | Mean | 95% CI |
| **Binary PA** | 1.00 | 1.00, 1.00 | 1.00 | 0.99, 1.00 | 1.00 | 0.99, 1.00 |
| **Continuous PA** | 0.00 | 0.00, 0.00 | 0.31 | 0.00, 0.98 | $3.51 \times 10^{-8}$ | $2.66 \times 10^{-16}$, $1.21 \times 10^{-7}$ |
| **OD** | 0.98 | 0.97, 0.99 | 0.65 | 0.21, 0.96 | 0.50 | 0.08, 0.82 |
| **qPCR** | 0.93 | 0.86, 0.97 | 0.98 | 0.95, 1.00 | 0.84 | 0.74, 0.92 |

**Supplementary Table 7: Estimates from the species only model for repeatability and heritability.** Estimates of repeatability are taken from model (2) and estimates of phylogenetic heritability (the variation explained by the host phylogeny) are taken from model (1). * indicates the phylogenetic heritability calculated as the proportion of variation that is attributed to phylogeny divided by the total variation (phylogenetic, non-phylogenetic between-strain, and within-strain variation) as opposed to phylogenetic and non-phylogenetic between-strain variation. PA = plaque assay, CI = credible interval.

| | **Repeatability** | | **Phylogenetic heritability** | | **Phylogenetic heritability of total variance*** | |
|---|---|---|---|---|---|---|
| **Method** | Mean | 95% CI | Mean | 95% CI | Mean | 95% CI |
| **Binary PA** | 1.00 | 1.00, 1.00 | 0.25 | 0.00, 0.91 | 0.25 | $3.18 \times 10^{-8}$, 0.91 |
| **Continuous PA** | 0.00 | 0.00, 0.00 | 0.38 | 0.00, 1.00 | $1.40 \times 10^{-7}$ | $2.22 \times 10^{-14}$, $3.95 \times 10^{-7}$ |
| **OD** | 0.83 | 0.69, 0.95 | 0.23 | 0.00, 0.81 | 0.17 | $1.70 \times 10^{-8}$, 0.66 |
| **qPCR** | 0.93 | 0.87, 0.97 | 0.28 | 0.00, 0.93 | 0.23 | $3.04 \times 10^{-7}$, 0.83 |

**Supplementary Table 8: Inter-strain correlations between methods for assessing host range in a within-*aureus* model.** Numbers show the mean estimates for the correlation strength (r, white cells) and slope (β, grey cells) between pairs of methods, with 95% credible intervals (CIs) indicated in brackets. The correlations and slopes were calculated with columns as x and rows as y. Estimates with CIs that do not span zero are highlighted in bold. PA = plaque assay, *value on a probit scale.

| | Binary PA | Continuous PA | OD | qPCR |
|---|---|---|---|---|
| **Binary PA** | - | - | -0.12 (-0.67, 0.48) | -0.67 (-0.39, 0.82) |
| **Continuous PA** | - | - | 0.02 (-0.94, 0.91) | 0.03 (-0.91, 0.95) |
| **OD** | **0.50* (0.49, 0.51)** | 0.03 (-0.81, 0.86) | - | **0.77 (0.47, 0.96)** |
| **qPCR** | **0.51* (0.49, 0.53)** | -0.24 (-2.19, 1.94) | **1.74 (0.53, 3.02)** | - |

**Supplementary Table 9: Inter-species correlations between methods for assessing host range in a between-species model.** Numbers show the mean estimates for the correlation strength (r, white cells) and slope (β, grey cells) between pairs of methods, with 95% credible intervals (CIs) indicated in brackets. The correlations and slopes were calculated with columns as x and rows as y. Estimates with CIs that do not span zero are highlighted in bold. PA = plaque assay, *value on a probit scale.

| | Binary PA | Continuous PA | OD | qPCR |
|---|---|---|---|---|
| **Binary PA** | - | - | **0.79 (0.46, 0.99)** | **0.86 (0.62, 1.00)** |
| **Continuous PA** | - | - | 0.03 (-0.96, 0.99) | 0.03 (-0.99, 0.98) |
| **OD** | **0.50* (0.50, 0.51)** | -0.00 (-0.09, 0.07) | - | **0.89 (0.72, 1.00)** |
| **qPCR** | **0.52* (0.50, 0.56)** | -0.02 (-0.57, 0.43) | **5.92 (3.47, 8.39)** | - |

**Table S20: *Staphylococcus* sequences used in the FastQ Screen to determine if all sequences used in this study belong to *Staphylococcus*/*S. aureus*.**

| Species | Strain | GenBank Accession Number |
|---|---|---|
| S. aureus | LGA251 | GCA_000237265.1 |
| S. caeli | 82B | GCA_900097965.1 |
| S. capitis | AYP1020 | GCA_001028645.1 |
| S. cohnii | SNUDS-2 | GCA_001990205.1 |
| S. condimenti | DSM_11674 | GCA_001922405.1 |
| S. edaphicus | CM_8730 | GCF_002614725.1 |
| S. epidermidis | ATCC_12228 | GCA_000007645.1 |
| S. haemolyticus | JCSC1435 | GCA_000009865.1 |
| S. hominis | K1 | GCA_002850375.1 |
| S. hyicus | ATCC_11249 | GCA_000816085.1 |
| S. kloosii | CNV2_Masurca_598 | GCA_001593625.1 |
| S. nepalensis | JS1 | GCA_002442895.1 |
| S. pasteuri | SP1 | GCA_000494875.1 |
| S. pettenkoferi | FDAARGOS_288 | GCA_002208805.2 |
| S. pseudintermedius | HKU10-03 | GCA_000185885.1 |
| S. saprophyticus | ATCC_15305 | GCA_007814115.1 |
| S. sciuri | FDAARGOS_285 | GCA_002209165.2 |
| S. simiae | NCTC13838 | GCA_900187055.1 |
| S. simulans | FDAARGOS_124 | GCA_001559115.2 |
| S. stepanovicii | NCTC13839 | GCA_900187075.1 |
| S. succinus | 14BME20 | GCF_001902315.1 |
| S. warneri | SG1 | GCA_007668085.1 |
| S. xylosus | HKUOPL8 | GCA_000706685.1 |

### 3. Supplementary Figures

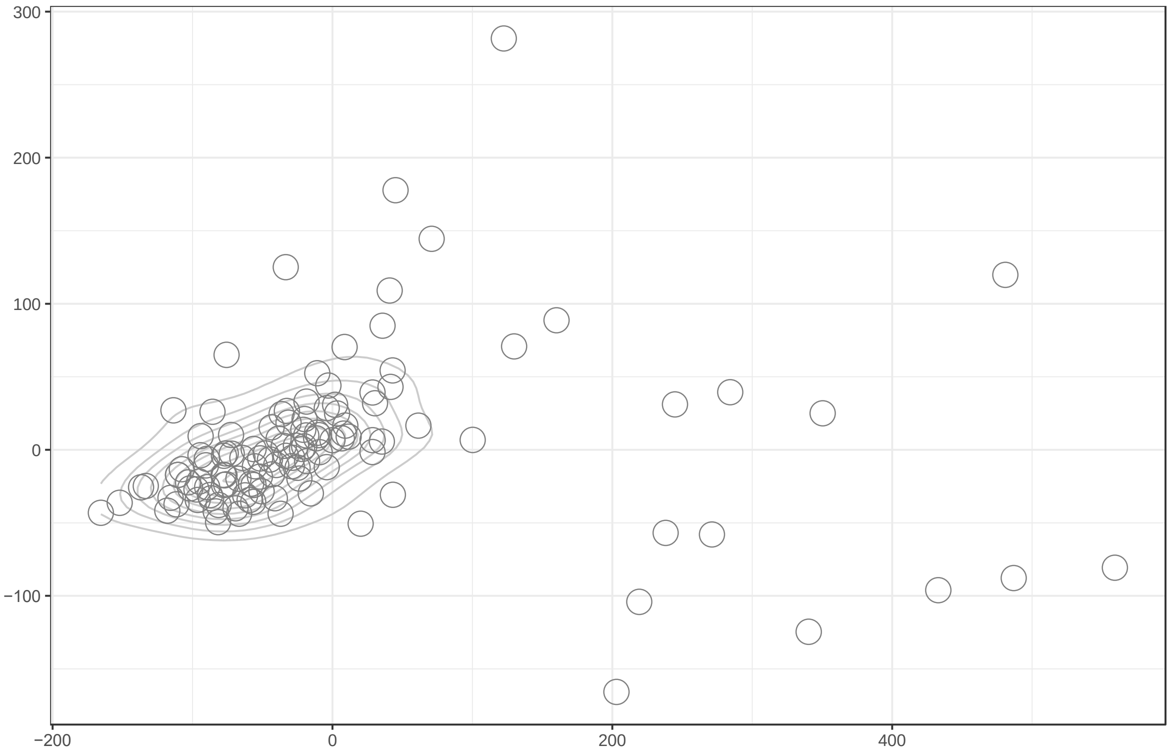


**Supplementary Figure 1: A comparison of all 123 trees on 64 tips using the Kendall Colijn metric vector (mid-point rooted gene trees).** MDS visualization of tree distances when $\lambda = 0$. As the projection often requires that multiple trees are plotted at the same co-ordinates, contour lines are used to indicate the density of points.

4. Supplementary References


1. Martin M. Cutadapt removes adapter sequences from high-throughput sequencing reads. EMBnet.journal. 2011 May;17(1):10.
2. Joshi N, Fass J. Sickle: A sliding-window, adaptive, quality-based trimming tool for FastQ files (Version 1.33) [Software]. Available at https://github.com/najoshi/sickle. 2011;2011.
3. Wingett S. FastQ Screen - Contamination screening for NGS data. http://www.Bioinformatics.Babraham.Ac.Uk/Projects/fastq-screen/. 2011.
4. Bankevich A, Nurk S, Antipov D, Gurevich AA, Dvorkin M, Kulikov AS, et al. SPAdes: a new genome assembly algorithm and its applications to single-cell sequencing. J Comput Biol. 2012 May;19(5):455–77.
5. Gurevich A, Saveliev V, Vyahhi N, Tesler G. QUAST: Quality assessment tool for genome assemblies. Bioinformatics. 2013 Apr;29(8):1072–5.
6. Langmead B, Salzberg SL. Fast gapped-read alignment with Bowtie 2. Nat Methods. 2012 Mar;9(4):357–9.
7. Wood DE, Salzberg SL. Kraken: Ultrafast metagenomic sequence classification using exact alignments. Genome Biol. 2014 Mar;15(3):1–12.
8. Page AJ, Taylor B, Keane JA. Multilocus sequence typing by blast from de novo assemblies against PubMLST. J Open Source Softw. 2016;1(8).
9. Schijffelen MJ, Boel CE, van Strijp JA, Fluit AC. Whole genome analysis of a livestock-associated methicillin-resistant Staphylococcus aureus ST398 isolate from a case of human endocarditis. BMC Genomics. 2010 Jun;11(1):376.
10. Frampton M, Houlston R. Generation of Artificial FASTQ Files to Evaluate the Performance of Next-Generation Sequencing Pipelines. Badger JH, editor. PLoS One. 2012 Nov;7(11):e49110.
11. Croucher NJ, Page AJ, Connor TR, Delaney AJ, Keane JA, Bentley SD, et al. Rapid phylogenetic analysis of large samples of recombinant bacterial whole genome sequences using Gubbins. Nucleic Acids Res. 2015 Feb;43(3):e15.
12. Didelot X, Wilson DJ. ClonalFrameML: Efficient Inference of Recombination in Whole Bacterial Genomes. PLoS Comput Biol. 2015;11(2):e1004041.
13. Price LB, Stegger M, Hasman H, Aziz M, Larsen J, Andersen PS, et al. *Staphylococcus aureus* CC398: Host adaptation and emergence of methicillin resistance in livestock. MBio. 2012 Mar;3(1):1–6.
14. Stamatakis A. RAxML version 8: a tool for phylogenetic analysis and post-analysis of large phylogenies. Bioinformatics [Internet]. 2014 May 1;30(9):1312–3. Available from: https://doi.org/10.1093/bioinformatics/btu033
15. Letunic I, Bork P. Interactive Tree Of Life (iTOL) v5: an online tool for phylogenetic tree display and annotation. Nucleic Acids Res [Internet]. 2021 Jul 2;49(W1):W293–6. Available from: https://doi.org/10.1093/nar/gkab301
16. Seemann T. Prokka: Rapid prokaryotic genome annotation. Bioinformatics. 2014 Jul;30(14):2068–9.
17. Tonkin-Hill G, MacAlasdair N, Ruis C, Weimann A, Horesh G, Lees JA, et al. Producing polished prokaryotic pangenomes with the Panaroo pipeline. Genome Biol. 2020 Jul;21(1):1–21.
18. Paradis E, Schliep K. ape 5.0: an environment for modern phylogenetics and evolutionary analyses in R. Bioinformatics [Internet]. 2019 Feb 1;35(3):526–8. Available from: https://doi.org/10.1093/bioinformatics/bty633

19. Jombart T, Kendall M, Almagro-Garcia J, Colijn C. treespace: Statistical exploration of landscapes of phylogenetic trees. Mol Ecol Resour. 2017 Nov;17(6):1385–92.

20. Kendall M, Colijn C. Mapping Phylogenetic Trees to Reveal Distinct Patterns of Evolution. Mol Biol Evol. 2016 Oct;33(10):2735–43.

21. Shapiro B, Rambaut A, Drummond AJ. Choosing appropriate substitution models for the phylogenetic analysis of protein-coding sequences. Vol. 23, Molecular biology and evolution. United States; 2006. p. 7–9.

22. Rambaut A, Suchard M, Xie D, Drummond A. Tracer v1.6 [Internet]. 2014. Available from: http://beast.bio.ed.ac.uk/Tracer
